# *Stenotrophomonas oleivorans sp. nov.* A polycyclic aromatic hydrocarbon-degrading strain isolated from crude oil contaminated soil

**DOI:** 10.64898/2026.01.18.700197

**Authors:** Temidayo O Elufisan, Isabel Cristina Rodríguez-Luna, Alejandro Sánchez-Varela, Patricia Bustos, Luis Lozano, Edgar Dantán Gonzalez, Omotayo Oyedara Opemipo, Jose Correa-Basurto, Alan Rubén Estrada-Pérez, Diana V. Cortés-Espinosa, Miguel Angel Villalobos-Lopez, Xianwu Guo

## Abstract

ASS1 was isolated as a motile Stenotrophomonas strain from crude oil-contaminated soils in Tabasco, Mexico. We characterized this strain using physiological and biochemical traits. ASS1 grew at temperature 25–37 (optimally at 37 ° C) and at pH 6 to 8 (optimally at pH 7 to 8). The assembled genome has a total length of 4.56MB with a G + C content of 66.6%. The 16S rRNA gene sequence analysis confirmed that this strain belongs to the genus Stenotrophomonas. Based on the 16S rRNA analysis, *Stenotrophomonas geniculata* ATCC 19374 is the closest species, and it shares 99.86% similarity with ASS1. Similarly, a phylogenomic tree based on core genome sequence revealed that the closest species to ASS1 is *Stenotrophomonas geniculata* ATCC 19374. The major fatty acids in ASS1 are C16:0, antesio C15:0, iso C12:0, iso C15:0, iso C17:0 and C18:0. The genome of ASS1 consists of 4,373,402 bp. The Average Nucleotide Identity (ANI) values for ASS1 which it shared with its closest phylogenetic neighbors, are *Stenotrophomonas geniculata* ATCC 19374 = JCM 13324 [T] 92.66 %, *Stenotrophomonas maltophilia* 13637[T] 92.15%, *Stenotrophomonas maltophilia* K279a 92.13% *Stenotrophomonas maltophilia* R551-3 92.15% *Stenotrophomonas maltophilia* MTCC 434 [T] 92.08% and *Pseudomonas hibisicicola* ATCC [T] 91.66%. ASS1 possesses genes that are essential for the degradation of Polycyclic Aromatic hydrocarbon. Genes such as 1, 2 dihydroxyl 1, 2 dihydronaphthalene dehydrogenase; MG068 17425, homologous to 2 hydroxyl chromene 2 carboxylate isomerases; MG 18055, homologous to salicylaldehyde dehydrogenase and MG068 20095, homologous to naphthalene 1, 2 dioxygenases were identified in ASS1. The dDDH value between ASS1 and its closest neighbor *Stenotrophomonas geniculata* ATCC 19374 = JCM 13324 [T] is 50%, which is the highest for all the typed species and as such we proposed that ASS1 is a novel species with the name *Stenotrophomonas oleivorans* sp. nov. sp. nov. and ASS1^T^ as the typed strain

## Introduction

*Stenotrophomonas* is a genus of a group of non-fermentative gram-negative bacteria in the class Gamma-proteobacteria (Ryan et al. 2009). The first member of the genus was isolated as *Bacterium bookeri* from the pleural fluid of a patient with oral carcinoma by Edward in 1943(Hugh and Ryschenkow 1961; Denton and Kerr 1998). Later *B. bookeri* and five previously identified *Pseudomonas alcaligenes* were reclassified as *Pseudomonas maltophilia* (Hugh and leifson 1963). In a related event, Komagata, et al., 1974 reviewed the classification of four *Pseudomonas melanogen*a strains and discovered that these strains were actually *Pseudomonas maltophilia* (Komagata et al. 1974). Similarly, several other studies have reviewed the genus resulting in the reclassification of some *Alcaligenes faecalis* as *Pseudomonas maltophilia*, and reclassification of *Pseudomonas maltophilia* as *Xanthomonas maltophilia*. These classifications were based on DNA-rRNA hybridization, analysis of 16S rRNA cistron, comparative enzymology, particularly the absence of NADP-linked dehydrogenases; the occurrence of the same type of ubiquinone in the cell membrane and cellular fatty acid composition (Ulrich and Needham 1953; Palleroni et al. 1973; Yang et al. 1993; Swings et al. 2009). The inability to adequately place *P. maltophilia* in the genus *Xanthomonas* using the 16S rRNA sequence made Palleroni and Bradbury propose a new genus with the name *Stenotrophomonas* for all the bacteria classified as Xanthomonas maltophilia by Swing et al., 2009 (Palleroni and Bradbury 1993). *Stenotrophomonas maltophilia* was the only member of the genus at that time. The reclassification was authenticated by Nesme et al., (Nesme et al. 1995) who used the restriction mapping of PCR amplified 16S rRNA gene to distinguish *Xanthomonas* and *Stenotrophomonas*

The name *Stenotrophomonas* was adopted as a description of the feeding pattern of *Stenotrophomonas maltophilia* (Stanier et al., 2009). *Stenotrophomonas* means a unit feeding on few substrates. It was originally believed that *Stenotrophomonas* strains were mainly associated with pathogenicity or diseases conditions because *S. maltophilia* was the only species in the genus then. However, several studies have led to the isolation of other non-pathogenic species from different environments (Geng et al. 2010). As a result of further isolation and characterization, the genus presently has 24 species, with 22 validly published name on the list of prokaryotic nomenclature (https://lpsn.dsmz.de/search?word=Stenotrophomonas) so far.

The genus has undergone various taxonomic revision after its recognition as a distinct genus: the transfer of *Stenotrophomonas dokdonensis* to the family *Pseudoxanthomonas dokdonensis* and the identification of *Stenotrophomonas africana* as a later synonym of the *S. maltophilia* are good examples of such taxonomic revision in the genus (Coenye 2004; Rasko et al. 2011). Patil et al., (2016) proposed a review of the genus following a pan genomic analysis of 19 sequenced genomes. They suggested that *S. maltophilia* and its related synonyms should be regrouped as *S. maltophila* complex (Smc). They reported that members of the *S. maltophilia* complex include *S. africana*, *Pseudomonas geniculata, Pseudomonas beteli, Stenotrophomonas pavanii and Stenotrophomonas sepilia* (Patil et al. 2016b; Gautam et al. 2021).

The advent of genome sequencing and analyses has revolutionized the taxonomy of bacteria. Genome sequencing and analysis provides a comprehensive perspective on the classification and delineation of bacteria species. Taxogenomics techniques have been employed in separating once cumbersome species groups into the appropriate categories (Patil et al. 2016a, 2018). Taxogenomics involves the use of complete genome sequence tools such as average nucleotide identity (ANI), digital DNA–DNA hybridization (dDDH), multi-locus sequence typing, phylogenetic reconstruction, and average amino acid identity for the classification of bacteria (Patil et al. 2016b; Oyedara et al. 2018; Elufisan et al. 2019).

The pan genome analysis of the genus *Stenotrophomonas* showed that the genus is still open and there are possibilities of finding or encountering new species (Patil et al. 2016b). In this article, we reported a new species of *Stenotrophomonas* isolated from crude oil contaminated soils during our study on the evolution and diversity of *Stenotrophomonas* species in the environment. ASS1 was recovered from crude-oil contaminated soil sample, and it grew effectively in naphthalene, phenanthrene and anthracene supplemented Bushnell Hass minimal medium respectively. Further genomic analysis of this strain also confirmed the presence of some genes like 1, 2 dihydroxyl 1, 2 dihydronaphthalene dehydrogenase; 2 hydroxyl chromene 2 carboxylate isomerases; salicylaldehyde dehydrogenase and to naphthalene 1, 2 dioxygenases which have been previously reported to be involved in the degradation of PAH(Das et al. 2015; Pal et al. 2017; Elufisan et al. 2019, 2020b)

## Method

### Isolation and growth condition

The bacterium in this study was isolated from crude oil contaminated soil recovered form Tabasco Mexico (17°52′26.9″N 92°29′12.4″W). The soil samples were aseptically collected into a sterile 50 ml conical tube and immediately transferred to the lab in sterile condition. 10 mg of the soil was transferred into 90 mL of sterile water and agitated at 150 rpm for 1 h to homogenize the soil sample. Once homogenized the mixture was centrifuged at 3000 rpm for 20 min. The supernatant from the soil sample was then used for serial dilution in distill water from a factor of 10^-1^ a factor of 10^-8^. The diluents were enriched in Luria broth and incubated at 30 °C for 24 h. The broths were then distributed in different dilutions between 10^-1^ to 10^-8^ for further inoculation on Stenotrophomonas Vancomycin Imipenem agar (SVIA) to select for *Stenotrophomonas* species. A 100µl of each dilution was spread on SVIA agar and incubated at 30 ° for the possible recovery of Stenotrophomonas species. Following these activities, yellow or green colonies which appeared on plates were characterized using biochemical, MALDI-TOF and molecular techniques.

### Microscopy

The morphology of the bacterial cells was determined by using Olympus compound light microscope and scanning electron microscope. The hanging drop method was used to evaluate the motility of the isolates as observed under the light microscope. For SEM analysis, bacterial cells were fixed with 8% paraformaldehyde (1:1) for 4 h at 4 °C. The cells were washed and dehydrated with graded ethanol (20%, 40%, 60%, 80%, 100%; each 20 min). They were placed inside the critical point dryer (Quorum, K850, UK) and dried with liquid CO_2_. After drying, they were stored in a desiccator until the image was checked with SEM microscope. The dried samples were gold coated for 30 seconds on a SPI model 12150-AB coater. All samples were analyzed with a scanning electron microscope.

### Phenotypic and biochemical characteristics of the isolates

The colonies appeared as yellow slightly raised on SVIA agar and Luria Bertani agar. The isolate appeared colorless on McConkey agar confirming that they are non-lactose fermenters. The biochemical characteristics of ASS1 were determined using the conventional method of biochemical characterization according to Bergey’s manual of determinative bacteriological studies (Palleroni and Bradbury 1993). A detailed description of the method for determining the biochemical characteristics and the antibiotic susceptibility studies for ASS1 has been reported in our previous article (Elufisan et al. 2020a).

### Chemotaxonomic characterization

Fatty acids from the bacterial strain were extracted as described by Brian & Gardner 1961 (Brian and Gardner 1967). Briefly, 4 loopfuls of bacterial colonies were harvested from an overnight grown culture in tryptic soy agar and transferred into 13 × 100 mL tubes to which 1 mL of saponification reagent was already added. The mixtures were vortexed vigorously for 10 seconds after which the tubes were heated in boiling water in the water bath for 5 min. After heating, the mixtures were once again vortexed vigorously for 10 seconds. And the tubes were again transferred into a water bath for further heating for 30 min. The mixtures were allowed to cool and followed by the addition of a 2 mL methylation reagent. The tubes were vortexed vigorously and heated in the water bath at 80° C for 10 min. To extract the methyl ester, 1.25 mL of extraction buffer was added to each mixture after cooling. The mix was then remixed by gentle tumbling and transferred for further analysis with the UHPLC.

The lipid analysis was carried out with UHPLC 1290 Infinity II (Agilent Technologies) with an Acquity UPLC BEH C18 2.1 × 50 mm, 1.7 µm column at 50 ± 0.5 °C (Waters). as described below. 10 µL of each sample were injected by triplicate using a flow of 0.3 mL/min and eluted with the following gradient conditions (A: 0.1 % formic acid in water and B: 0.1 % formic acid in methanol): 55 % at 0 min, 90 % B at 3 min, 45 % B at 10 min and maintained for 2 min. Mass spectra were acquired in a Q-TOF 6545A system (Agilent Technologies) with Agilent Mass Hunter LC/MS Data Acquisition Software Version B.08.00 Build 8.00.8058.3 SP1 and using an Agilent Dual AJS ESI ion source operated in positive polarity. Source conditions were set as follows: sheath gas flow and temperature of 10 L/min and 350 °C, respectively, nebulizer pressure of 40 (pounds per square inch (psi)), drying gas flow and temperature of 10 L/min 320 °C, respectively, capillary voltage 3500 V, nozzle voltage of 1000 V and 175 V for the fragmentor. Q-TOF instrument was operated in 2GHz extended dynamic range mode. MS spectra were acquired at a rate of 1 spectrum/s in a 100 – 3000 m/z range. Internal reference mass calibration ions used during data acquisition were: 121.050873 and 922.009798.

### LC-ESI-MS/MS Data Processing

All data acquired were first inspected using Agilent Mass Hunter Qualitative Analysis Software B.07.00 Build 7.7.7024.29 SP2 to find spectra corresponding to fatty acid compounds previously reported (Sánchez-Castro et al. 2017) using the Find by Formula (FBF) algorithm. Relative abundances of compounds found were retrieved from data and are shown supplementary table 1.

## Genomic analysis

### Strain identification and phylogenomic Investigation

Extracted DNAs from ASS1 previously identified by 16S rRNA partial gene (Elufisan et al. 2020a), were sequenced. The identity of the sequence was confirmed by blast search analysis on the NCBI database and the ezbiocloud database. The obtained sequences were then aligned with other *Stenotrophomonas*’ sequences retrieved from NCBI database on the MEGA X software. The aligned sequences were then used for phylogenetic reconstruction on MEGA X software. Similarly, an MLST tree based on 82 concatenated multi-locus sequence genes in *Stenotrophomonas* was drawn using the online autoMLST at https://automlst.ziemertlab.com. Similarly, bacterial pan-core genome analysis was carried out with bacterial pan genome analysis tool (BPGA) to generate a concatenated core genes sequence. The concatenated core genes were aligned using the muscles algorithm on MEGA XI and a core genome based-phylogenetic tree was drawn from the alignment using the neighbor joining algorithm with a bootstrap repeat of 1000. Presence of contamination in the genome was checked on https://www.ezbiocloud.net/tools/contest16s website (Lee et al. 2017).

### Strain identification with genome-based phylogeny and Taxonomy

The complete genome was sequenced with Illumina MiSeq and the obtained reads were assembled into contigs with SPAdes (version 3.12.0) (Bankevich et al. 2012). The G + C content for the genome was estimated from the assembly. The genome completeness was checked with CHECKM on the Kbase website (Allen et al. 2022). The functional genes and potential coding region were predicted with Prokka (version) (Seemann 2014) while the functional gene and potential coding region for the genome version available online was predicted with the NCBI Prokaryotic Genome Annotation Pipeline (PGAP)(Tatusova et al. 2016). Circular genome map was drawn with Proksee (Stothard and Wishart 2005). Pangenome analysis was carried out using bacteria pangenome analysis tool (BPGA) version 1.3 (Chaudhari et al., 2016) using the default parameter. The core, accessory, and unique genes in ASS1 were annotated with eggnog-mapper online server using the default parameter. The function of the fitness curve was determined with the formula f(x)= a*x^-b^ where a= 3478.54 and b= 0.485217.

The relatedness of the whole genome sequence with other members of the genus was evaluated using average nucleotide identity (ANI) value and digital DNA-DNA Hybridization (dDDH). The ANI was calculated using the JSspeciesWS (Richter et al. 2016) and the dDDH was estimated with the online GGDC server (version 3.0)(Meier-Kolthoff et al. 2013). To determine the novelty of the strain or species, we used the TYGS server for identifying type strains. TYGS server uses both 16S rRNA and dDDH for the identification of new species (Meier-Kolthoff and Göker 2019). Deep sequence analysis to identify the genes associated with PAH utilization was done on the JGI-IMG platform (Mavromatis et al. 2009) and antiSMASH v 6 (Blin et al. 2021) was used to predict the biosynthetic cluster in ASS1.

### Growth and survival in polycyclic aromatic hydrocarbon

ASS1’s ability to grow and survive in oil-contaminated sites was evaluated in a tolerance test with varying concentration of seven different PAHs (anthracene, anthraquinone, biphenyl, naphthalene, phenanthridine, phenanthrene and xylene) in a Bush Nell Hass medium. Each setup contains one of the seven hydrocarbons as the only carbon source in the Bushnell Hass medium. A final concentration of 5 mg/mL and 1 mg/mL was used to test bacterial growth and survival in PAHs in the tolerance assay. The hydrocarbon tolerance contains 1% of 100mg/ml of either naphthalene, phenanthridine, anthraquinones, biphenyl, phenanthrene and 1% of 40mg/ml of anthracene added to a in Bushnell Hass (BH) as the sole carbon respectively. This process was repeated with 5% of 100mg/ml of naphthalene, phenanthridine, anthraquinones, biphenyl, phenanthrene and 40 mg/ml with respect to anthracene to obtain a 5mg/ml final concentration. All hydrocarbons were dissolved in Dimethyl chloride and the solvent was left to evaporate before introducing the hydrocarbons into each culture medium. The experimental design has three setups which include the test experiment and two controls. All experiments were duplicates. A 100 µL of overnight grown culture of ASS1 washed in phosphate buffer was inoculated in 100 mL of each BH medium containing either 1% and 5% each of the above-mentioned PAH at a concentration of 100 mg/mL or 40mg/ml for anthracene in 250 mL Erlenmeyer flask while an uninoculated BH medium with hydrocarbon only and a BH medium with ASS1 only served as controls. All experimental set up were incubated at 30 ^°^C in a rotatory incubator at 200 rpm for 25 days following inoculation with test bacteria. ASS1’s tolerance was checked every two days using colony counting method. 100 µL of each serially diluted (10^-4^) inoculated medium was spread on Luria Bertani agar plates and were incubated at 30 ^°^C for 24 h after which the colony formed were counted. Spectrophotometric analysis was also carried out on culture from all experimental setup to corroborate the observations from colony counting method.

## Result and Discussion

### Physiological and chemotaxonomic characteristics

ASS1 appears as a long rod under the scanning electron microscope as shown in Figure 1. Motility assay showed that they are motile as they exhibit movement under the light microscope though the microscopic image did not show any flagella.

**Figure 1:**
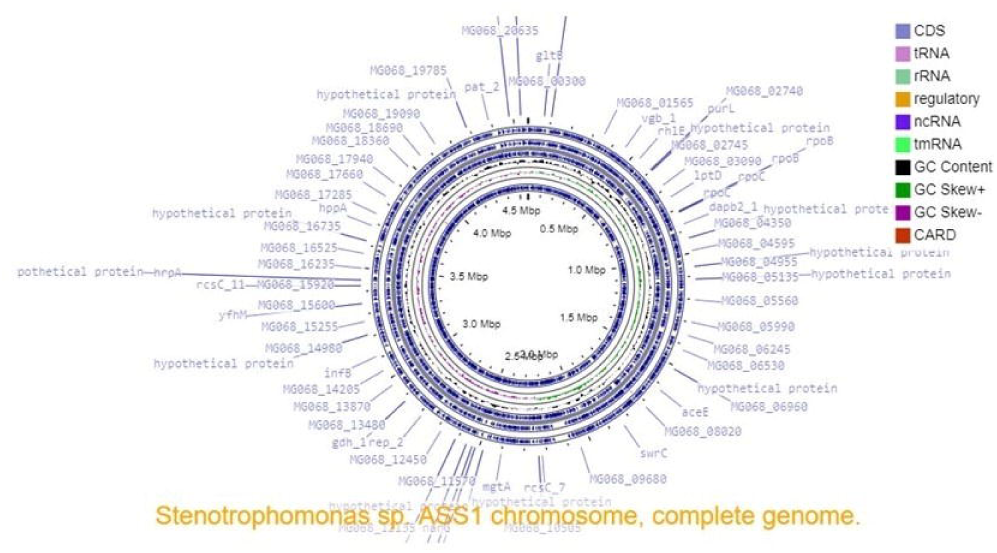
Phylogeny drawn from the alignment of the Stenotrophomonas 16S rRNA sequences of the typed strain and ASS1. The phylogeny was drawn using the neighbor-joining algorithm on MEGA XI with a bootstrap of 1000 repeats. ASS1 is in in yellow font while the outgroup strain in red font.

The biochemical characteristics of the isolates showed that ASS1 is a gram-negative non-lactose fermenter, positive for catalase production and negative for the oxidase activity. The biochemical characteristics of ASS1 compared with the available reports for the other published species are shown in Table 1. ASS1 differs from its closest species by not being able to hydrolyze gelatin whereas *Pseudomonas hibiscicola* and *Pseudomonas geniculata* from which it branched on the phylogenetic can do so. However, it could use galactose as a carbon source for growth whereas the two closer strains; *Pseudomonas hibiscicola* and *Pseudomonas geniculata* cannot.

Previous study have shown the possibility of using LC-MS (UHPLC-ESI-QTOF-MS) for identifying bacterial lipid profile (Řezanka et al. 2012). LC-MS can be used to detect the genetic differences among different strains based on their lipid content that may be associated with their metabolism, and growth. (https://pubmed.ncbi.nlm.nih.gov/29341867/). The analysis of the fatty acids for *Stenotrophomonas* sp. ASS1 showed that the dominant fatty acids are the C_16:0,_ iso-C_12:0_, iso-C_16:0_, iso-C_17:1w_9c, and C_18:0,_ C_18:1w_9C and C_19,_ making it different from others, as shown in Table S1. ASS1 differs significantly from the two closer species in that it lacks the C_10:0_, Iso C_11:0_, iso C_11:0_3-OH fatty acids, as well as Iso C_13:0_ 3-OH, C_13:0_ 2-OH, antesio-C15:0, C_15:0_ that are present in *P. geniculata* and *P. hibiscicola*

ASS1 displayed resistant to sulfamethoxazole-trimethoprim, chloramphenicol, imipenem, ceftriaxone, ceftazidime, ampicillin, doxycycline, gentamicin, and nitrofurantoin. It is however susceptible to levofloxacin and ofloxacin (Elufisan et al. 2020a). ASS1 grew effectively in the tested PAHs showing the best growth in anthracene (figure S5).

## Genomic analysis

### Strain identification and phylogenomic Investigation

The strain ASS1 was recovered from crude oil contaminated soil samples. The complete 16S rRNA fragment of ASS1 was retrieved from its genome and deposited in the GenBank with the accession number <u>ON323668</u>. ASS1 shares between 74%-99.86% similarity with other members of the genus *Stenotrophomonas* and *Lysobacter* which was used as the outgroup strain in the phylogenetic tree. The closest species to ASS1 is *Pseudomonas geniculata* 19374^T^ with which it shared 99.86% 16S rRNA identity (Figure 2). Some studies have shown that 16S rRNA similarity above 99% did not signify that the bacteria are the same species though it suggests it could be so (Bosshard et al. 2006; Mignard and Flandrois 2006; Janda and Abbott 2007; Caudill and Brayton 2022). Thus, we further analyzed it in-depth at genomic levels using dDDH and average nucleotide identity blast (ANIb) analysis, core genome-based phylogenetic analysis and MLST based phylogeny. the phylogeny drawn based on the core and complete genome sequence, which revealed that the closest species to ASS1 is the *Stenotrophomonas* (*Pseudomonas*) *geniculata* ATCC 19374^T,^ because they branched out from the same clade (Figure S1). The MLST based concatenated tree showed that ASS1 branched out from the *S. maltophilia* clade (Figure S2). All the genomic comparison values showed that ASS1 belongs to a new species.

**Figure 2:**
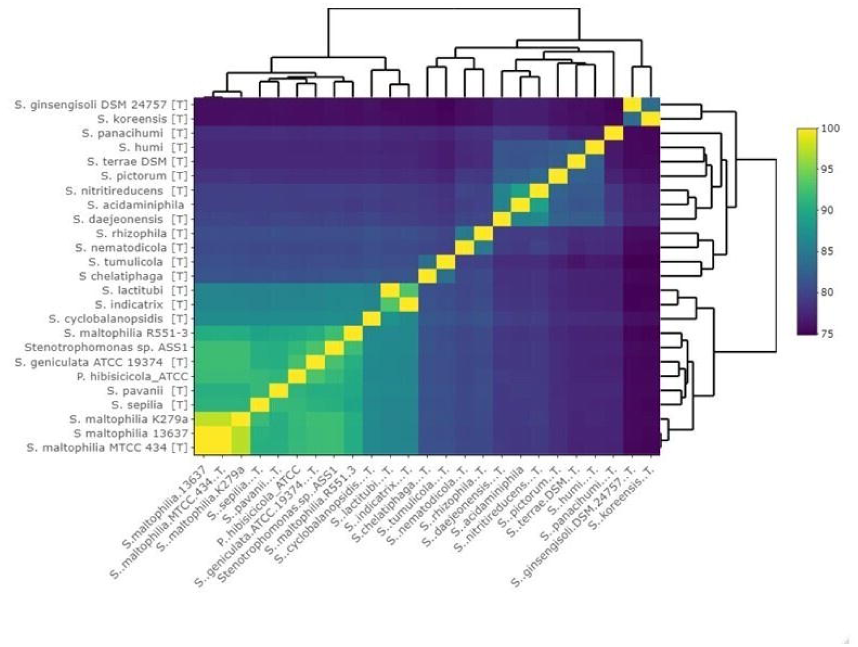
Scanning electron microscopic image of Stenotrophomonas sp. ASS1.

### Strain identification with genome-based phylogeny and Taxonomy

The genomic characteristics for ASS1 are shown in Table S2. The genomes consist of 4564481 bp. ASS1 was assembled to one contig each with a G + C value of 66.59. The pan-core genome analysis of ASS1 and 24 other *Stenotrophomonas* species retrieved from the NCBI data showed that it shared 1177 core genes (Figure 3). ASS1 possesses 2674 accessory genes and 130 unique genes. The α value for the fitness curve is 0.485217, which is much less than 1, indicating that the pangenome is open. So many unique genes and open pangenome indicate that more novel strains and new species could be found in the genus in the future.

**Figure 3:**
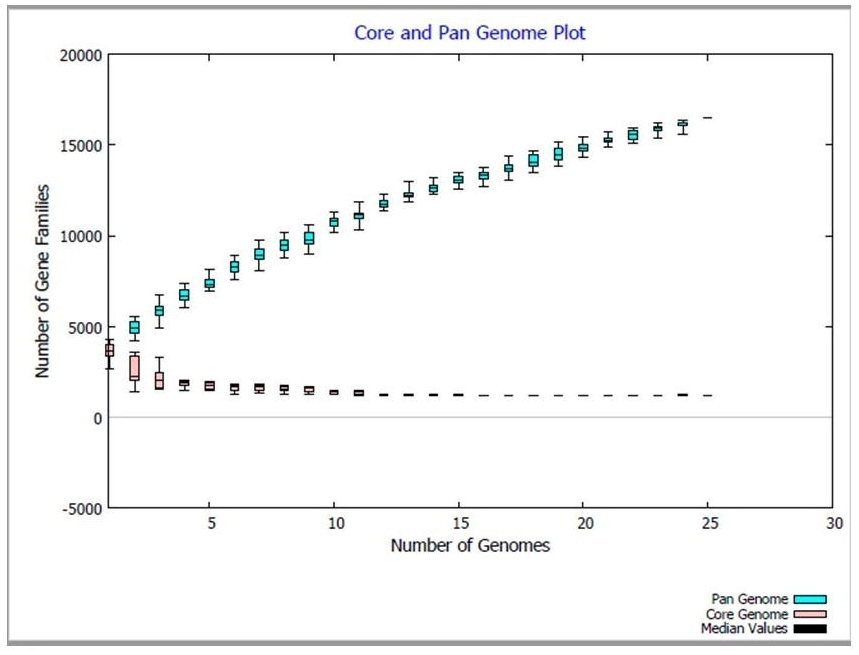
Core-Pan genome plot for ASS1 and other Stenotrophomonas species.

The annotation of the genes in the core, accessory and unique category showed that the genes that were associated with the degradation of hydrocarbon are distributed across them. The KEGG analysis showed that there are 44 genes in the core genome, 190 genes in the accessory genome and three unique genes of ASS1 are directly associated with the metabolism of various hydrocarbon (Table S3, S4, S5). ASS1 possesses several genes that are essential for the degradation of polycyclic aromatic hydrocarbon (PAH) such as MG06_03320, a gene that is homologous to 1, 2 dihydroxyl 1, 2 dihydronaphthalene dehydrogenase; MG068_17425, homologous to 2 hydroxyl chromene 2 carboxylate isomerases; MG_18055, homologous to salicylaldehyde dehydrogenase and MG068_20095, homologous to naphthalene 1, 2 dioxygenases. The presence of these genes in its genome explains why ASS1 grew and survives in an oil-contaminated site. These genes have been previously reported to be involved in the utilization and degradation of naphthalene by *Stenotrophomonas* and other bacteria (Das et al. 2015; Pal et al. 2017; Elufisan et al. 2020). In our previous study, we identified metabolites that were associated with the activities of some of the genes identified in ASS1 (Elufisan et al. 2020b).

6 biosynthetic clusters were found in ASS1 (Figure S4). The biosynthetic clusters include Arylpolyene cluster, 2 bacteriocin clusters, Lantipeptid**e** cluster, lassopeptide cluster, and Nrps Cluster. One of these clusters, Nrps is also associated with the degradation of hydrocarbon. The circular genome map of ASS1 is shown in figure 5.

The average nucleotide identity for *Stenotrophomonas* sp. ASS1 as shared with its closest genomes which are *Stenotrophomonas geniculata* ATCC 19374 = JCM 13324 [T] *Stenotrophomonas maltophilia* 13637[T] *Stenotrophomonas maltophilia* K279a *Stenotrophomonas maltophilia* R551-3 *Stenotrophomonas maltophilia* MTCC 434 [T] *and Pseudomonas hibiscicola* ATCC [T] are 92.66%, 921.15%, 92.13%, 92.15%, 92.08%, and 91.66% respectively (Figure 4). The choice of *Stenotrophomonas maltophilia* K279a *Stenotrophomonas maltophilia* R551-3 *Stenotrophomonas maltophilia* MTCC 434 in the analysis is due to the relatedness as revealed by ezbiocloud 16S rRNA similarity. The ANI value as estimated using ANIb on JSspeciesW is apparently below the threshold for classifying 2 bacteria as the same species, as a result we proposed ASS1 as a novel species in the genus *Stenotrophomonas* and as the type-strain. Analysis of ASS1 genome using the GGDC tool on the DSMZ website also confirmed that ASS1 is a novel species as the estimated dDDH value (50% against the closest species *Pseudomonas geniculata*) is below 70% with all the type strains of *Stenotrophomona*s species (Table 3). Similarly, TYGS tool (https://tygs.dsmz.de/) also identified *ASS1* as a potential new species.

**Figure 4:**
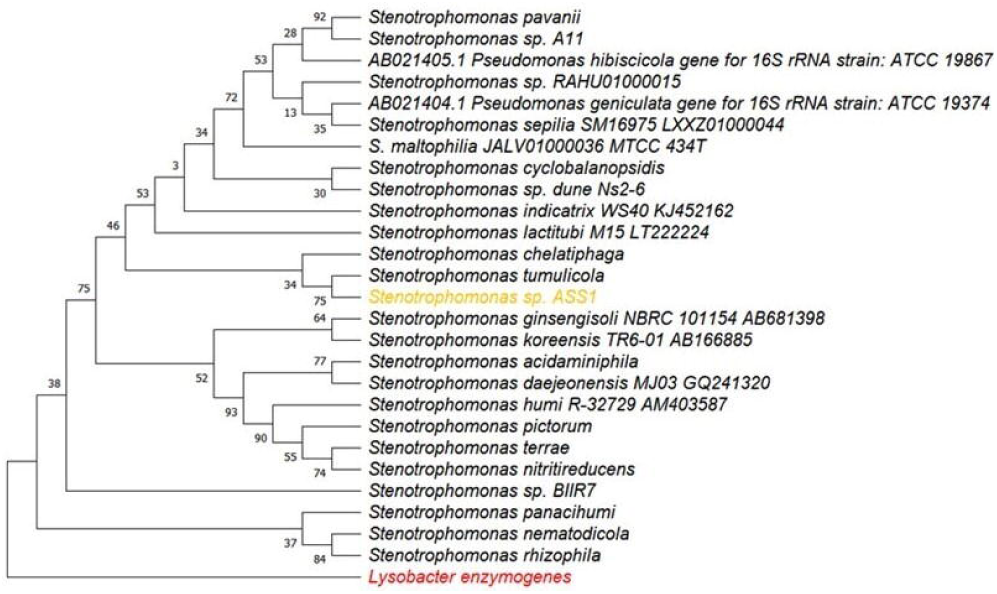
Heatmap showing the comparison of the average nucleotide identity of Stenotrophomonas oleivorans ASS1T.

**Figure 5:**
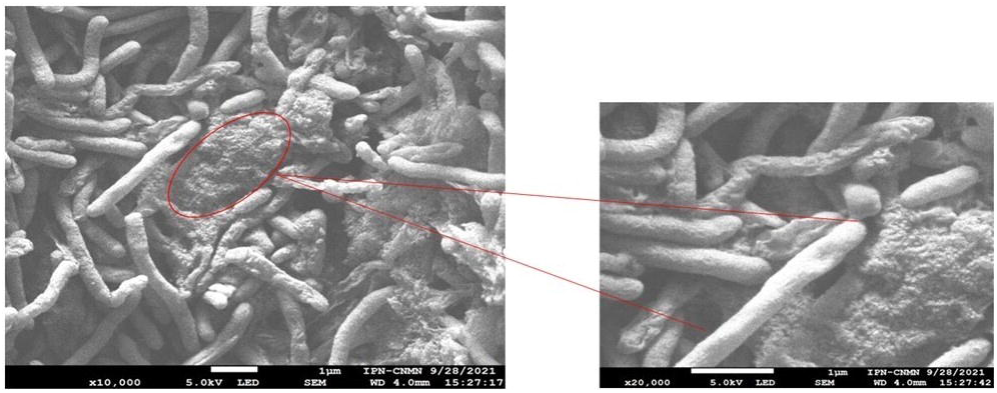
Stenotrophomonas sp. ASS1’ s growth on different PAHs determined by colony counts on every other day for 20 days.

**Figure 6:** Circular genome map of Stenotrophomonas sp ASS1.

### Taxonomic conclusion

Based on the distinct phylogenetic analyses within the currently available *Stenotrophomonas* species, biochemical characteristics and the low-level of DNA-DNA relatedness with the closely related species, it can be concluded that *Stenotrophomonas* sp. ASS1 is a novel species in the genus *Stenotrophomonas*. We therefore proposed the name *Stenotrophomonas oleivorans sp. nov*.

### Description of Stenotrophomonas oleivorans sp. nov

*Stenotrophomonas oleivorans sp. nov.* ASS1 (o.le.i.vo’rans. L. n. oleum, oil; L. part. adj. vorans, eating; N.L. fem. adj. oleivorans, eating oil, referring to…) is a gram-negative motile bacterium, which grew effectively at pH 7-8 and between 25-37 °C, and it grew most effectively at 37 °C after 48 h of incubation. *Stenotrophomonas petrorepungnans* ASS1 is positive for catalase production, but negative to the production of oxidase. It cannot hydrolyze gelatin, but hydrolyze starch, tween 80, and aesculin. It can use arabinose, glucose, fructose, mannose, and galactose as carbon sources for growth. It is a non-lactose fermenter and can utilize lysine and citrate for growth (Table 1). The incubation period is between 24 – 48 h on Luria agar and *Stenotrophomonas* Vancomycin Imipenem and Amphotericin agar, but the optimal incubation period is 48 hours on the predominant fatty-acids in the membrane of *Stenotrophomonas oleivorans sp. nov.* are the C_16:0,_ iso-C_12:0_, iso-C_16:0_, iso-C_17:1w_9c, and C_18:0,_ C_18:1w_9C and C_19,_ (Table S1).

ASS1 is resistant to sulfamethoxazole-trimethoprim, chloramphenicol, imipenem, ceftriaxone, ceftazidime, ampicillin, doxycycline, gentamicin, and nitrofurantoin. It is however susceptible to levofloxacin and ofloxacin (Elufisan et al. 2020).

ASS1 grew effectively in the tested PAHs and showed the best growth in anthracene.

The typed strain ASS1 was isolated from crude contaminated soil Villa Hermosa, Tabasco, Mexico (17°59′13″N 92°55′10″O w).

### Isolate Deposit information

The isolate has been deposited with the world culture collection center in Mexico with the name *Stenotrophomonas* sp. ASS1 WFCC 1006/CM-CNRG 934.

The complete genome sequence of *Stenotrophomonas oleivorans sp. nov* ASS1 has been deposited on NCBI database with the accession number <u>CP031167</u>

## Acknowledgment

We appreciate the support of PhD. Hugo Martínez Gutiérrez and PhD. María de Jesús Perea Flores from the Centro de Nanociencias y Micro y Nanotecnologías of Instituto Politécnico Nacional for the photo of scanned electron microscopy.

We want to thank Prof. Dr. rer. nat. Bernhard Schink for helping with the bacteria nomenclature.

## Authors’ contributions

Temidayo Oluyomi Elufisan; Perform the experiment and wrote the manuscript.

Isabel Cristina Luna-Rodrigruez participated in the design of the research project and provided some material for the experiment.

Alejandro Sanchez-Varela participated in the design of the experiment and participated in some experimental works. Patricia Bustos and Luiz Lozano helped with the genome assembly, annotation, and analysis.

Edgan Datan-Gozalez helped with the sequencing of the genome.

Omotayo Oyedara opemipo participated in the editing and review of the manuscript.

Jose Correa-Basurto did lipid analysis and participated in the writing and review of the manuscript. Alan Rubén Estrada Pérez did lipid analysis and participated in the writing and review of the manuscript.

Diana V Cortés-Espinosa helped with electron microscopy, data editing, and manuscript editing. Miguel-Angel Villalobos-Lopez, designed, supervised the research, and participated in the writing and editing of the manuscript.

Xianwu Guo designed, supervised the research, and participated in the writing and editing of the manuscript.

