## Supplementary file for "*Stenotrophomonas oleivorans sp. nov.* A polycyclic aromatic hydrocarbon-degrading strain isolated from crude oil contaminated soil"

| Table S1: Fatty acid methyl ester profile for ASS1 obtained from LC-MS. | | | | | | |  |  |
| --- | --- | --- | --- | --- | --- | --- | --- | --- |
| **Fatty acid** | **1** | **2** | **3** | **4** | **5** | **6** | **7** | **8** |
| **C_10:0_** | - | - | 0.6 | 0.6 | 0.9 | 2.0 | 0.89 | 0.87 |
| **Iso C_11:0_** | - | 3.7 | 3.0 | 3.3 | 3.0 | 4.2 | 2.56 | 2.64 |
| **iso C_11:0_ 3-OH** | - | 2.0 | 1.6 | 1.4 | 1.4 | 2.9 | 1.32 | 0.75 |
| **iso-C_12:0_ 3-OH** | 1.2 | - | 0.4 | - | - | 1.1 | 3.38 | 1.46 |
| **Iso- C_15:0_** | 3.7 | 23.8 | 17.9 | 30.1 | 29.4 | 10.6 | 26.19 | 26.85 |
| **Iso-C_13:0_ 3-OH** | - | 3.2 | 1.8 | 3.4 | 3.3 | 1.9 | 1.94 | 1.01 |
| **C_13:0_ 2-OH** | - | 1.7 | 0.9 | 0.9 | - | 1.6 | 0.4 | 0.3 |
| **Iso-C_15:1_** | - | 1.1 | 1.6 | - | - | 6.3 | 0.91 | 0.62 |
| **Anteiso-C_15:0_** | - | 19.4 | 22.2 | 23.3 | 13.3 | 10.8 | 11.62 | 16.60 |
| **C_15:0_** | - | 0.9 | 0.7 | - | - | 1.4 | - | - |
| **iso-C_16:1_** | 3.1 | - | - | - | - | - | - | - |
| **C_16:0_** | 10.2 | 9.4 | 8.6 | 7.0 | 13.7 | 7.4 | 12.88 | 7.29 |
| **C_16:1_** | 3.1 | - | - | - | - | - | - | - |
| **Iso-C_16:0_ 3-OH** | 7.3 | - | - | - | - | - | - | - |
| **C_16:1ɯ_9c** | 0.9 | 2.3 | 3.2 | 2.2 | 4.2 | 3.9 | 5.15 | 4.59 |
| **Iso-C_17:1ɯ_9c** | 2.8 | 7.5 | 10.0 | 4.0 | 3.9 | 1.5 | - | - |
| **Iso-C_17:0_** | - | 2.8 | 4.8 | 4.2 | 4.7 | - | - | - |
| **anteiso-C_17:0_** | - | - | 0.9 | 1.0 | - | - | 0.3 | 0.77 |
| **C_17:1ɯ_8C** | - | 0.7 | - | - | - | - | 0.24 | 0.24 |
| **C_18:0_** | 10.8 | - | - | - | - | - | 0.61 | 0.59 |
| **C_18:1ɯ_9_C_** | 39.9 | - | 1.0 | 1.7 | 1.2 | - | 1.88 | 2.37 |
| **C_19:1_** | 2.8 | - | - | - | - | - | - | - |

Strains: 1, ASS1, 2, *S. bentholitica* BII-R7T; 3, *S. rhizophila* DSM 14405T; 4, *S. pavanii* DSM 25135T; 5, *S. maltophilia* DSM 50170T; 6, *S. chelatiphaga* DSM 21508T; 7, *P. geniculata* JCM 13324T; 8. *P. hibiscicola* JCM 13361^T^.

_ means Not detected.

Data 2-6 were obtained from Sanchez-Castro et al., 2016 (Sánchez-Castro et al. 2017b)

And Data 7 and 8 were obtained from Gautam et al., 2021.

TableS2: List of MLST genes used for the concatenation

| TIGR00055 | uppS | Cell envelope | di-trans,poly-cis-decaprenylcistransferase |
| --- | --- | --- | --- |
| TIGR01850 | argC | Amino acid biosynthesis | N-acetyl-gamma-glutamyl-phosphate reductase |
| TIGR01959 | nuoF_fam | Energy metabolism | NADH oxidoreductase (quinone), F subunit |
| TIGR00060 | L18_bact | Protein synthesis | ribosomal protein uL18 |
| TIGR00138 | rsmG_gidB | Protein synthesis | 16S rRNA (guanine(527)-N(7))-methyltransferase RsmG |
| TIGR01978 | sufC | Biosynthesis of cofactors, prosthetic groups, and carriers | FeS assembly ATPase SufC |
| TIGR01164 | rplP_bact | Protein synthesis | ribosomal protein uL16 |
| TIGR00936 | ahcY | Energy metabolism | adenosylhomocysteinase |
| TIGR00033 | aroC | Amino acid biosynthesis | chorismate synthase |
| TIGR01302 | IMP_dehydrog | Purines, pyrimidines, nucleosides, and nucleotides | inosine-5'-monophosphate dehydrogenase |
| TIGR01135 | glmS | Central intermediary metabolism | glutamine-fructose-6-phosphate transaminase (isomerizing) |
| TIGR03723 | T6A_TsaD_YgjD | Protein synthesis | tRNA threonylcarbamoyl adenosine modification protein TsaD |
| TIGR01632 | L11_bact | Protein synthesis | ribosomal protein uL11 |
| TIGR01083 | nth | DNA metabolism | endonuclease III |
| TIGR00431 | TruB | Protein synthesis | tRNA pseudouridine(55) synthase |
| TIGR02970 | succ_dehyd_cytB | Energy metabolism | succinate dehydrogenase, cytochrome b556 subunit |
| TIGR02027 | rpoA | Transcription | DNA-directed RNA polymerase, alpha subunit |
| TIGR00959 | ffh | Protein fate | signal recognition particle protein |
| TIGR00337 | PyrG | Purines, pyrimidines, nucleosides, and nucleotides | CTP synthase |
| TIGR00928 | purB | Purines, pyrimidines, nucleosides, and nucleotides | adenylosuccinate lyase |
| TIGR01207 | rmlA | Cell envelope | glucose-1-phosphate thymidylyltransferase |
| TIGR01163 | rpe | Energy metabolism | ribulose-phosphate 3-epimerase |
| TIGR00510 | lipA | Biosynthesis of cofactors, prosthetic groups, and carriers | lipoyl synthase |
| TIGR00487 | IF-2 | Protein synthesis | translation initiation factor IF-2 |
| TIGR00628 | ung | DNA metabolism | uracil-DNA glycosylase |
| TIGR00962 | atpA | Energy metabolism | ATP synthase F1, alpha subunit |
| TIGR02891 | CtaD_CoxA | Energy metabolism | cytochrome c oxidase, subunit I |
| TIGR01816 | sdhA_forward | Energy metabolism | succinate dehydrogenase, flavoprotein subunit |
| TIGR01063 | gyrA | DNA metabolism | DNA gyrase, A subunit |
| TIGR01009 | rpsC_bact | Protein synthesis | ribosomal protein uS3 |
| TIGR01021 | rpsE_bact | Protein synthesis | ribosomal protein uS5 |
| TIGR01049 | rpsJ_bact | Protein synthesis | ribosomal protein uS10 |
| TIGR01341 | aconitase_1 | Energy metabolism | aconitate hydratase 1 |
| TIGR01024 | rplS_bact | Protein synthesis | ribosomal protein bL19 |
| TIGR00168 | infC | Protein synthesis | translation initiation factor IF-3 |
| TIGR01134 | purF | Purines, pyrimidines, nucleosides, and nucleotides | amidophosphoribosyltransferase |
| TIGR00041 | DTMP_kinase | Purines, pyrimidines, nucleosides, and nucleotides | dTMP kinase |
| TIGR01966 | RNasePH | Transcription | ribonuclease PH |
| TIGR00382 | clpX | Protein fate | ATP-dependent Clp protease, ATP-binding subunit ClpX |
| TIGR00244 | TIGR00244 | Regulatory functions | transcriptional regulator NrdR |
| TIGR00065 | ftsZ | Cellular processes | cell division protein FtsZ |
| TIGR00048 | rRNA_mod_RlmN | Protein synthesis | 23S rRNA (adenine(2503)-C(2))-methyltransferase |
| TIGR00165 | S18 | Protein synthesis | ribosomal protein bS18 |
| TIGR00063 | folE | Biosynthesis of cofactors, prosthetic groups, and carriers | GTP cyclohydrolase I |
| TIGR02729 | Obg_CgtA | Protein synthesis | Obg family GTPase CgtA |
| TIGR02075 | pyrH_bact | Purines, pyrimidines, nucleosides, and nucleotides | UMP kinase |
| TIGR00580 | mfd | DNA metabolism | transcription-repair coupling factor |
| TIGR00228 | ruvC | DNA metabolism | crossover junction endodeoxyribonuclease RuvC |
| TIGR00459 | aspS_bact | Protein synthesis | aspartate--tRNA ligase |
| TIGR00635 | ruvB | DNA metabolism | Holliday junction DNA helicase RuvB |
| TIGR03632 | uS11_bact | Protein synthesis | ribosomal protein uS11 |
| TIGR00223 | panD | Biosynthesis of cofactors, prosthetic groups, and carriers | aspartate 1-decarboxylase |
| TIGR03594 | GTPase_EngA | Protein synthesis | ribosome-associated GTPase EngA |
| TIGR03346 | chaperone_ClpB | Protein fate | ATP-dependent chaperone protein ClpB |
| TIGR00418 | thrS | Protein synthesis | threonine--tRNA ligase |
| TIGR03263 | guanyl_kin | Purines, pyrimidines, nucleosides, and nucleotides | guanylate kinase |
| TIGR02350 | prok_dnaK | Protein fate | chaperone protein DnaK |
| TIGR00612 | ispG_gcpE | Biosynthesis of cofactors, prosthetic groups, and carriers | 4-hydroxy-3-methylbut-2-en-1-yl diphosphate synthase |
| TIGR00184 | purA | Purines, pyrimidines, nucleosides, and nucleotides | adenylosuccinate synthase |
| TIGR01059 | gyrB | DNA metabolism | DNA gyrase, B subunit |
| TIGR02013 | rpoB | Transcription | DNA-directed RNA polymerase, beta subunit |
| TIGR00713 | hemL | Biosynthesis of cofactors, prosthetic groups, and carriers | glutamate-1-semialdehyde-2,1-aminomutase |
| TIGR00499 | lysS_bact | Protein synthesis | lysine--tRNA ligase |
| TIGR02274 | dCTP_deam | Purines, pyrimidines, nucleosides, and nucleotides | deoxycytidine triphosphate deaminase |
| TIGR00615 | recR | DNA metabolism | recombination protein RecR |
| TIGR00362 | DnaA | DNA metabolism | chromosomal replication initiator protein DnaA |
| TIGR01980 | sufB | Biosynthesis of cofactors, prosthetic groups, and carriers | FeS assembly protein SufB |
| TIGR00243 | Dxr | Biosynthesis of cofactors, prosthetic groups, and carriers | 1-deoxy-D-xylulose 5-phosphate reductoisomerase |
| TIGR00088 | trmD | Protein synthesis | tRNA (guanine(37)-N(1))-methyltransferase |
| TIGR00263 | trpB | Amino acid biosynthesis | tryptophan synthase, beta subunit |
| TIGR01171 | rplB_bact | Protein synthesis | ribosomal protein uL2 |
| TIGR01099 | galU | Cell envelope | UTP--glucose-1-phosphate uridylyltransferase |
| TIGR00493 | clpP | Protein fate | ATP-dependent Clp endopeptidase, proteolytic subunit ClpP |
| TIGR01039 | atpD | Energy metabolism | ATP synthase F1, beta subunit |
| TIGR01071 | rplO_bact | Protein synthesis | ribosomal protein uL15 |
| TIGR01029 | rpsG_bact | Protein synthesis | ribosomal protein uS7 |
| TIGR00482 | TIGR00482 | Biosynthesis of cofactors, prosthetic groups, and carriers | nicotinate (nicotinamide) nucleotide adenylyltransferase |
| TIGR00521 | coaBC_dfp | Biosynthesis of cofactors, prosthetic groups, and carriers | phosphopantothenoylcysteine decarboxylase / phosphopantothenate--cysteine ligase |
| TIGR01051 | topA_bact | DNA metabolism | DNA topoisomerase I |
| TIGR01011 | rpsB_bact | Protein synthesis | ribosomal protein uS2 |
| TIGR01296 | asd_B | Amino acid biosynthesis | aspartate-semialdehyde dehydrogenase |

TableS4: Identified hydrocarbon degrading genes in the core genome of ***Stenotrophomonas petrorepugnans*** ASS1

| query | seed_ortholog | eggNOG_OGs | COG_category | Description | Preferred_name | EC | KEGG_ko |
| --- | --- | --- | --- | --- | --- | --- | --- |
| core/91/2/WP_132810464.1 | 522373.Smlt3167 | COG2902@1\|root,COG2902@2\|Bacteria,1MXNV@1224\|Proteobacteria,1RQVZ@1236\|Gammaproteobacteria,1X46R@135614\|Xanthomonadales | E | glutamate dehydrogenase | - | 1.4.1.2 | ko:K15371 |
| core/289/2/WP_032128799.1 | 522373.Smlt2219 | COG0458@1\|root,COG0458@2\|Bacteria,1MUDZ@1224\|Proteobacteria,1RPIU@1236\|Gammaproteobacteria,1X3MZ@135614\|Xanthomonadales | F | Four CarB-CarA dimers form the carbamoyl phosphate synthetase holoenzyme that catalyzes the production of carbamoyl phosphate | carB | 6.3.5.5 | ko:K01955 |
| core/580/2/WP_132811103.1 | 522373.Smlt0778 | COG2352@1\|root,COG2352@2\|Bacteria,1MUD5@1224\|Proteobacteria,1RPTP@1236\|Gammaproteobacteria,1X4MI@135614\|Xanthomonadales | C | Forms oxaloacetate, a four-carbon dicarboxylic acid source for the tricarboxylic acid cycle | ppc | 4.1.1.31 | ko:K01595 |
| core/860/2/WP_132810381.1 | 391008.Smal_2422 | COG0574@1\|root,COG1080@1\|root,COG0574@2\|Bacteria,COG1080@2\|Bacteria,1MU0R@1224\|Proteobacteria,1RP3T@1236\|Gammaproteobacteria,1X4C5@135614\|Xanthomonadales | G | Catalyzes the phosphorylation of pyruvate to phosphoenolpyruvate | ppsA | 2.7.9.2 | ko:K01007 |
| core/1135/2/WP_132810818.1 | 522373.Smlt4120 | COG1249@1\|root,COG1249@2\|Bacteria,1MU2U@1224\|Proteobacteria,1RMFF@1236\|Gammaproteobacteria,1X3MV@135614\|Xanthomonadales | C | dehydrogenase | lpdA | 1.8.1.4 | ko:K00382 |
| core/1266/2/WP_132810596.1 | 522373.Smlt3556 | COG1024@1\|root,COG1250@1\|root,COG1024@2\|Bacteria,COG1250@2\|Bacteria,1MU9P@1224\|Proteobacteria,1RMZ8@1236\|Gammaproteobacteria,1X486@135614\|Xanthomonadales | I | 3-hydroxyacyl-coa dehydrogenase | fadJ | 1.1.1.35,4.2.1.17,5.1.2.3 | ko:K01782 |
| core/1303/2/WP_132808808.1 | 522373.Smlt0237 | COG4770@1\|root,COG4770@2\|Bacteria,1P6RE@1224\|Proteobacteria,1RM95@1236\|Gammaproteobacteria,1X2ZC@135614\|Xanthomonadales | I | carboxylase | - | 6.4.1.4 | ko:K01968 |
| core/1325/2/WP_132811074.1 | 391008.Smal_4013 | COG0507@1\|root,COG0507@2\|Bacteria,1MW43@1224\|Proteobacteria,1RPA0@1236\|Gammaproteobacteria,1X3QZ@135614\|Xanthomonadales | L | A helicase nuclease that prepares dsDNA breaks (DSB) for recombinational DNA repair. Binds to DSBs and unwinds DNA via a highly rapid and processive ATP-dependent bidirectional helicase activity. Unwinds dsDNA until it encounters a Chi (crossover hotspot instigator) sequence from the 3' direction. Cuts ssDNA a few nucleotides 3' to the Chi site. The properties and activities of the enzyme are changed at Chi. The Chi-altered holoenzyme produces a long 3'-ssDNA overhang and facilitates RecA-binding to the ssDNA for homologous DNA recombination and repair. Holoenzyme degrades any linearized DNA that is unable to undergo homologous recombination. In the holoenzyme this subunit has ssDNA-dependent ATPase and 5'-3' helicase activity. When added to pre-assembled RecBC greatly stimulates nuclease activity and augments holoenzyme processivity. Negatively regulates the RecA-loading ability of RecBCD | recD | 3.1.11.5 | ko:K03581 |
| core/1426/2/WP_032128463.1 | 522373.Smlt4623 | COG0365@1\|root,COG0365@2\|Bacteria,1MUF5@1224\|Proteobacteria,1RMNZ@1236\|Gammaproteobacteria,1X34U@135614\|Xanthomonadales | I | Catalyzes the conversion of acetate into acetyl-CoA (AcCoA), an essential intermediate at the junction of anabolic and catabolic pathways. AcsA undergoes a two-step reaction. In the first half reaction, AcsA combines acetate with ATP to form acetyl-adenylate (AcAMP) intermediate. In the second half reaction, it can then transfer the acetyl group from AcAMP to the sulfhydryl group of CoA, forming the product AcCoA | acsA | 6.2.1.1 | ko:K01895 |
| core/1482/2/WP_049462121.1 | 522373.Smlt3355 | COG1154@1\|root,COG1154@2\|Bacteria,1MUSJ@1224\|Proteobacteria,1RNQD@1236\|Gammaproteobacteria,1X2YR@135614\|Xanthomonadales | H | Catalyzes the acyloin condensation reaction between C atoms 2 and 3 of pyruvate and glyceraldehyde 3-phosphate to yield 1-deoxy-D-xylulose-5-phosphate (DXP) | dxs | 2.2.1.7 | ko:K01662 |
| core/1599/2/WP_132810686.1 | 391008.Smal_3188 | COG3240@1\|root,COG5571@1\|root,COG3240@2\|Bacteria,COG5571@2\|Bacteria,1MWDI@1224\|Proteobacteria,1S2RQ@1236\|Gammaproteobacteria,1X3QI@135614\|Xanthomonadales | IN | esterase | estA | - | ko:K12686 |
| core/1719/2/WP_071229676.1 | 391008.Smal_2777 | COG1960@1\|root,COG1960@2\|Bacteria,1MU20@1224\|Proteobacteria,1RNV1@1236\|Gammaproteobacteria,1X364@135614\|Xanthomonadales | I | acyl-coa dehydrogenase | - | - | - |
| core/1721/2/WP_132809817.1 | 391008.Smal_1537 | COG1053@1\|root,COG1053@2\|Bacteria,1MU5M@1224\|Proteobacteria,1RMU2@1236\|Gammaproteobacteria,1X4JC@135614\|Xanthomonadales | C | Belongs to the FAD-dependent oxidoreductase 2 family. FRD SDH subfamily | sdhA | 1.3.5.1,1.3.5.4 | ko:K00239 |
| core/1861/2/WP_107431288.1 | 391008.Smal_2890 | COG1785@1\|root,COG1785@2\|Bacteria,1MXI2@1224\|Proteobacteria,1RNG8@1236\|Gammaproteobacteria,1X4AG@135614\|Xanthomonadales | P | Belongs to the alkaline phosphatase family | phoA | 3.1.3.1 | ko:K01077 |
| core/1862/2/WP_132809283.1 | 391008.Smal_0720 | COG3914@1\|root,COG4235@1\|root,COG3914@2\|Bacteria,COG4235@2\|Bacteria,1QU3U@1224\|Proteobacteria,1T2GT@1236\|Gammaproteobacteria,1XD7Y@135614\|Xanthomonadales | O | Glycosyl transferase family 41 | - | - | - |
| core/2105/2/WP_049400582.1 | 391008.Smal_0197 | COG4799@1\|root,COG4799@2\|Bacteria,1MVAX@1224\|Proteobacteria,1RNV5@1236\|Gammaproteobacteria,1X4BD@135614\|Xanthomonadales | I | Acetyl-CoA carboxylase, carboxyltransferase component (subunits alpha and beta) | - | 6.4.1.4 | ko:K01969 |
| core/2133/2/WP_132809268.1 | 391008.Smal_0688 | COG3634@1\|root,COG3634@2\|Bacteria,1MUKD@1224\|Proteobacteria,1RNC7@1236\|Gammaproteobacteria,1X4HZ@135614\|Xanthomonadales | O | Alkyl hydroperoxide reductase | ahpF | - | ko:K03387 |
| core/2154/2/WP_071229725.1 | 522373.Smlt3282 | COG1271@1\|root,COG1271@2\|Bacteria,1MV60@1224\|Proteobacteria,1RN2U@1236\|Gammaproteobacteria,1X4FJ@135614\|Xanthomonadales | C | oxidase, subunit | cydA | 1.10.3.14 | ko:K00425 |
| core/2313/2/WP_132809980.1 | 522373.Smlt2132 | COG1012@1\|root,COG1012@2\|Bacteria,1MW72@1224\|Proteobacteria,1RY9G@1236\|Gammaproteobacteria,1X3B2@135614\|Xanthomonadales | C | belongs to the aldehyde dehydrogenase family | - | 1.2.1.3 | ko:K00128 |
| core/2998/2/WP_132809698.1 | 391008.Smal_1258 | COG0750@1\|root,COG0750@2\|Bacteria,1MU91@1224\|Proteobacteria,1RMIX@1236\|Gammaproteobacteria,1X3KG@135614\|Xanthomonadales | M | zinc metalloprotease | rseP | - | ko:K11749 |
| core/3058/2/WP_014037873.1 | 522373.Smlt3315 | COG0402@1\|root,COG0402@2\|Bacteria,1MVPA@1224\|Proteobacteria,1RN13@1236\|Gammaproteobacteria,1X44W@135614\|Xanthomonadales | F | Catalyzes the hydrolytic cleavage of a carbon-halogen bond in N-ethylammeline | - | 3.5.4.28,3.5.4.31 | ko:K12960 |
| core/3256/2/WP_132810921.1 | 522373.Smlt4329 | COG3508@1\|root,COG3508@2\|Bacteria,1MV9G@1224\|Proteobacteria,1RQG2@1236\|Gammaproteobacteria,1X40V@135614\|Xanthomonadales | Q | Involved in the catabolism of homogentisate (2,5- dihydroxyphenylacetate or 2,5-OH-PhAc), a central intermediate in the degradation of phenylalanine and tyrosine. Catalyzes the oxidative ring cleavage of the aromatic ring of homogentisate to yield maleylacetoacetate | hmgA | 1.13.11.5 | ko:K00451 |
| core/3294/2/WP_240792140.1 | 391008.Smal_1338 | COG0665@1\|root,COG0665@2\|Bacteria,1MVGP@1224\|Proteobacteria,1RNJ9@1236\|Gammaproteobacteria,1X30K@135614\|Xanthomonadales | E | Oxidoreductase | - | - | ko:K09471 |
| core/3455/2/WP_132810594.1 | 522373.Smlt3534 | COG0654@1\|root,COG0654@2\|Bacteria,1MUZP@1224\|Proteobacteria,1S37X@1236\|Gammaproteobacteria,1X2Z7@135614\|Xanthomonadales | CH | Oxidoreductase | - | - | - |
| core/3470/2/WP_132809259.1 | 522373.Smlt0825 | COG0654@1\|root,COG0654@2\|Bacteria,1MU6I@1224\|Proteobacteria,1RMS3@1236\|Gammaproteobacteria,1X3RQ@135614\|Xanthomonadales | CH | hydroxylase | ubiH | - | ko:K03185 |
| core/3618/2/WP_181453208.1 | 522373.Smlt3097 | COG0077@1\|root,COG1605@1\|root,COG0077@2\|Bacteria,COG1605@2\|Bacteria,1MU60@1224\|Proteobacteria,1RNRD@1236\|Gammaproteobacteria,1X4R7@135614\|Xanthomonadales | E | Prephenate dehydratase | pheA | 4.2.1.51,5.4.99.5 | ko:K14170 |
| core/3889/2/WP_126929264.1 | 522373.Smlt4687 | COG1092@1\|root,COG1092@2\|Bacteria,1MUGB@1224\|Proteobacteria,1RN7Z@1236\|Gammaproteobacteria,1X40U@135614\|Xanthomonadales | J | Oxidoreductase | - | 2.1.1.191 | ko:K06969 |
| core/3930/2/WP_049463155.1 | 522373.Smlt3174 | COG1960@1\|root,COG1960@2\|Bacteria,1MUDR@1224\|Proteobacteria,1RMMJ@1236\|Gammaproteobacteria,1X2XI@135614\|Xanthomonadales | I | acyl-CoA dehydrogenase | acdA | - | - |
| core/3986/2/WP_132809261.1 | 391008.Smal_0676 | COG0654@1\|root,COG0654@2\|Bacteria,1MU6I@1224\|Proteobacteria,1RND5@1236\|Gammaproteobacteria,1X3DS@135614\|Xanthomonadales | CH | Catalyzes the formation of 2-octaprenyl-3-methyl-5-hydroxy-6-methoxy-1,4-benzoquinol from 2-octaprenyl-3-methyl-6-methoxy-1,4-benzoquinol | visC | - | ko:K03184,ko:K18800 |
| core/4060/2/WP_071228793.1 | 522373.Smlt0199 | COG1960@1\|root,COG1960@2\|Bacteria,1MUK0@1224\|Proteobacteria,1RNBX@1236\|Gammaproteobacteria,1X45S@135614\|Xanthomonadales | I | Acyl-CoA dehydrogenase | gcdH | 1.3.8.6 | ko:K00252 |
| core/4144/2/WP_071229128.1 | 522373.Smlt3429 | COG0082@1\|root,COG0082@2\|Bacteria,1MU98@1224\|Proteobacteria,1RMQS@1236\|Gammaproteobacteria,1X365@135614\|Xanthomonadales | E | Catalyzes the anti-1,4-elimination of the C-3 phosphate and the C-6 proR hydrogen from 5-enolpyruvylshikimate-3-phosphate (EPSP) to yield chorismate, which is the branch point compound that serves as the starting substrate for the three terminal pathways of aromatic amino acid biosynthesis. This reaction introduces a second double bond into the aromatic ring system | aroC | 4.2.3.5 | ko:K01736 |
| core/4380/2/WP_132809820.1 | 391008.Smal_1545 | COG1194@1\|root,COG1194@2\|Bacteria,1MUD4@1224\|Proteobacteria,1RMBT@1236\|Gammaproteobacteria,1X4F8@135614\|Xanthomonadales | L | glycosylase | mutY | - | ko:K03575 |
| core/4475/2/WP_132810771.1 | 391008.Smal_3387 | COG1062@1\|root,COG1062@2\|Bacteria,1MUK4@1224\|Proteobacteria,1RNQ4@1236\|Gammaproteobacteria,1X3UT@135614\|Xanthomonadales | C | Belongs to the zinc-containing alcohol dehydrogenase family. Class-III subfamily | frmA | 1.1.1.1,1.1.1.284 | ko:K00121 |
| core/4596/2/WP_132809263.1 | 391008.Smal_0678 | COG2933@1\|root,COG2933@2\|Bacteria,1MWBM@1224\|Proteobacteria,1RMSB@1236\|Gammaproteobacteria,1X494@135614\|Xanthomonadales | J | Belongs to the class I-like SAM-binding methyltransferase superfamily. RNA methyltransferase RlmE family. RlmM subfamily | rlmM | 2.1.1.186 | ko:K06968 |
| core/4616/2/WP_049401686.1 | 522373.Smlt4338 | COG1071@1\|root,COG1071@2\|Bacteria,1MU5R@1224\|Proteobacteria,1RREX@1236\|Gammaproteobacteria,1X3WI@135614\|Xanthomonadales | C | COG1071 Pyruvate 2-oxoglutarate dehydrogenase complex, dehydrogenase (E1) component, eukaryotic type, alpha subunit | pdhA | 1.2.4.1 | ko:K00161 |
| core/5011/2/WP_005407597.1 | 391008.Smal_0136 | COG0240@1\|root,COG0240@2\|Bacteria,1MUU3@1224\|Proteobacteria,1RPQ7@1236\|Gammaproteobacteria,1X2XQ@135614\|Xanthomonadales | I | Glycerol-3-phosphate dehydrogenase | gpsA | 1.1.1.94 | ko:K00057 |
| core/5064/2/WP_132809309.1 | 522373.Smlt0954 | COG0604@1\|root,COG0604@2\|Bacteria,1MU4N@1224\|Proteobacteria,1RNSV@1236\|Gammaproteobacteria,1X3D6@135614\|Xanthomonadales | C | Belongs to the zinc-containing alcohol dehydrogenase family. Quinone oxidoreductase subfamily | - | 1.6.5.5 | ko:K00344 |
| core/5356/2/WP_132809096.1 | 391008.Smal_0481 | COG0179@1\|root,COG0179@2\|Bacteria,1MV0V@1224\|Proteobacteria,1RNYV@1236\|Gammaproteobacteria,1X4ZT@135614\|Xanthomonadales | Q | 2-keto-4-pentenoate hydratase | uptA | 3.7.1.2 | ko:K16171 |
| core/5678/2/WP_049424359.1 | 1429851.X548_04285 | COG0761@1\|root,COG0761@2\|Bacteria,1MU7G@1224\|Proteobacteria,1RMN8@1236\|Gammaproteobacteria,1X4RS@135614\|Xanthomonadales | IM | Catalyzes the conversion of 1-hydroxy-2-methyl-2-(E)- butenyl 4-diphosphate (HMBPP) into a mixture of isopentenyl diphosphate (IPP) and dimethylallyl diphosphate (DMAPP). Acts in the terminal step of the DOXP MEP pathway for isoprenoid precursor biosynthesis | ispH | 1.17.7.4 | ko:K03527 |
| core/7635/2/WP_019337645.1 | 391008.Smal_0278 | COG4221@1\|root,COG4221@2\|Bacteria,1MUF8@1224\|Proteobacteria,1RMKM@1236\|Gammaproteobacteria,1X40Q@135614\|Xanthomonadales | S | Belongs to the short-chain dehydrogenases reductases (SDR) family | - | 1.1.1.276,1.1.1.381 | ko:K05886,ko:K16066 |
| core/7705/2/WP_032128162.1 | 522373.Smlt0374 | COG1011@1\|root,COG1011@2\|Bacteria,1N0I6@1224\|Proteobacteria,1RQ41@1236\|Gammaproteobacteria,1X45A@135614\|Xanthomonadales | S | Hydrolase | - | 3.1.3.102,3.1.3.104 | ko:K20862 |
| core/7767/2/WP_032130254.1 | 391008.Smal_2750 | COG1028@1\|root,COG1028@2\|Bacteria,1MUEV@1224\|Proteobacteria,1RNH2@1236\|Gammaproteobacteria,1X4C7@135614\|Xanthomonadales | IQ | Belongs to the short-chain dehydrogenases reductases (SDR) family | phbB | 1.1.1.36 | ko:K00023 |
| core/8491/2/WP_132810826.1 | 522373.Smlt4132 | COG0400@1\|root,COG0400@2\|Bacteria,1RA02@1224\|Proteobacteria,1S24F@1236\|Gammaproteobacteria,1X4YF@135614\|Xanthomonadales | S | Carboxylesterase | - | - | ko:K06999 |
| core/8546/2/WP_132811077.1 | 522373.Smlt4673 | COG2854@1\|root,COG2854@2\|Bacteria,1N022@1224\|Proteobacteria,1TA6U@1236\|Gammaproteobacteria,1X35G@135614\|Xanthomonadales | Q | toluene tolerance | yrbC | - | ko:K07323 |
| core/11673/2/WP_071229045.1 | 391008.Smal_3123 | COG0824@1\|root,COG0824@2\|Bacteria,1MZH6@1224\|Proteobacteria,1S93F@1236\|Gammaproteobacteria,1X6W5@135614\|Xanthomonadales | S | Acyl-CoA thioesterase | - | - | ko:K07107 |
| core/11759/2/WP_049423471.1 | 522373.Smlt4239 | COG0757@1\|root,COG0757@2\|Bacteria,1RDDT@1224\|Proteobacteria,1S3PX@1236\|Gammaproteobacteria,1X6EW@135614\|Xanthomonadales | E | Catalyzes a trans-dehydration via an enolate intermediate | aroQ | 4.2.1.10 | ko:K03786 |
| core/13977/2/WP_132809451.1 | 391008.Smal_0994 | COG2146@1\|root,COG2146@2\|Bacteria,1N8PE@1224\|Proteobacteria,1SG29@1236\|Gammaproteobacteria,1X6VK@135614\|Xanthomonadales | P | Benzene 1,2-dioxygenase | bedB | - | ko:K05710 |

Table S5 Identified hydrocarbon degrading related genes in the unique genome of ***Stenotrophomonas* *petrorepugnans***

| query | seed_ortholog | evalue | eggNOG_OGs | COG_category | Description | Preferred_name |
| --- | --- | --- | --- | --- | --- | --- |
| unique/4426/2/WP_132809988.1 | 1537917.JU82_00910 | 7.38e-159 | COG0438@1\|root,COG0438@2\|Bacteria,1MVA7@1224\|Proteobacteria,42P3H@68525\|delta/epsilon subdivisions,2YMIR@29547\|Epsilonproteobacteria | M | glycosyl transferase group 1 | - |
| unique/4977/2/WP_132810171.1 | 59538.XP_005976350.1 | 3.63e-228 | COG0604@1\|root,KOG1198@2759\|Eukaryota,39EFV@33154\|Opisthokonta,3C0GH@33208\|Metazoa,3DGSB@33213\|Bilateria | C | Zinc-binding dehydrogenase | - |
| unique/5048/2/WP_132809986.1 | 981336.F944_01150 | 8.84e-85 | COG0438@1\|root,COG0438@2\|Bacteria,1N9EV@1224\|Proteobacteria,1RYRV@1236\|Gammaproteobacteria,3NPSW@468\|Moraxellaceae | M | Glycosyl transferases group 1 | rfbU |

Table S6: Identified genes that are associated with hydrocarbon degradation in the accessory genome of ***Stenotrophomonas* *petrorepugnans***

| query | seed_ortholog | evalue | eggNOG_OGs | max_annot_lvl | COG_category | Description | Preferred_name | EC | KEGG_ko |
| --- | --- | --- | --- | --- | --- | --- | --- | --- | --- |
| accessory/231/2/WP_132810759.1 | 522373.Smlt3935 | 0.0 | COG3537@1\|root,COG3537@2\|Bacteria,1N86N@1224\|Proteobacteria,1RZCC@1236\|Gammaproteobacteria,1X4Z0@135614\|Xanthomonadales | 135614\|Xanthomonadales | G | Glycosyl hydrolase family 92 | - | - | - |
| accessory/257/2/WP_132810969.1 | 391008.Smal_3814 | 0.0 | COG1501@1\|root,COG1501@2\|Bacteria,1MWNJ@1224\|Proteobacteria,1RMJ9@1236\|Gammaproteobacteria,1X3WR@135614\|Xanthomonadales | 135614\|Xanthomonadales | G | Belongs to the glycosyl hydrolase 31 family | - | - | - |
| accessory/331/2/WP_132810013.1 | 522373.Smlt2179 | 0.0 | COG1629@1\|root,COG1629@2\|Bacteria,COG4771@2\|Bacteria,1MV8W@1224\|Proteobacteria,1RMQA@1236\|Gammaproteobacteria,1X4VK@135614\|Xanthomonadales | 135614\|Xanthomonadales | P | TonB-dependent receptor | - | - | - |
| accessory/355/2/WP_121504397.1 | 340.xcc-b100_2957 | 0.0 | COG1629@1\|root,COG1629@2\|Bacteria,COG4771@2\|Bacteria,1N7IE@1224\|Proteobacteria,1RQ90@1236\|Gammaproteobacteria,1XC9Y@135614\|Xanthomonadales | 135614\|Xanthomonadales | P | TonB dependent receptor | - | - | - |
| accessory/371/2/WP_132809721.1 | 522373.Smlt1542 | 0.0 | COG2199@1\|root,COG3292@1\|root,COG3292@2\|Bacteria,COG3706@2\|Bacteria,1R9NB@1224\|Proteobacteria,1S1MC@1236\|Gammaproteobacteria,1XCAK@135614\|Xanthomonadales | 135614\|Xanthomonadales | T | Two component regulator propeller | - | - | - |
| accessory/383/2/WP_132810968.1 | 522373.Smlt4432 | 0.0 | COG1629@1\|root,COG4771@2\|Bacteria,1MU9K@1224\|Proteobacteria,1RMTG@1236\|Gammaproteobacteria,1XA00@135614\|Xanthomonadales | 135614\|Xanthomonadales | P | TonB dependent receptor | - | - | - |
| accessory/400/2/WP_132810348.1 | 522373.Smlt2919 | 0.0 | COG1472@1\|root,COG1472@2\|Bacteria,1MVIV@1224\|Proteobacteria,1RMA0@1236\|Gammaproteobacteria,1X4BZ@135614\|Xanthomonadales | 135614\|Xanthomonadales | G | Belongs to the glycosyl hydrolase 3 family | bglS | 3.2.1.21 | ko:K05349 |
| accessory/471/2/WP_132810163.1 | 391008.Smal_2065 | 0.0 | COG1629@1\|root,COG1629@2\|Bacteria,COG4771@2\|Bacteria,1NAZC@1224\|Proteobacteria,1S063@1236\|Gammaproteobacteria,1X9P4@135614\|Xanthomonadales | 135614\|Xanthomonadales | P | TonB dependent receptor | - | - | - |
| accessory/486/2/WP_132810557.1 | 1429851.X548_12950 | 0.0 | COG1629@1\|root,COG1629@2\|Bacteria,COG4771@2\|Bacteria,1MU9K@1224\|Proteobacteria,1RMTG@1236\|Gammaproteobacteria,1X4HX@135614\|Xanthomonadales | 135614\|Xanthomonadales | P | TonB-dependent receptor | - | - | - |
| accessory/531/2/WP_132809291.1 | 391008.Smal_0739 | 0.0 | COG1629@1\|root,COG1629@2\|Bacteria,COG4771@2\|Bacteria,1MU9K@1224\|Proteobacteria,1RMTG@1236\|Gammaproteobacteria,1X4KS@135614\|Xanthomonadales | 135614\|Xanthomonadales | P | TonB dependent receptor | - | - | - |
| accessory/549/2/WP_132810045.1 | 522373.Smlt2241 | 0.0 | COG1048@1\|root,COG1048@2\|Bacteria,1MU9T@1224\|Proteobacteria,1RN5I@1236\|Gammaproteobacteria,1X3SP@135614\|Xanthomonadales | 135614\|Xanthomonadales | C | Catalyzes the isomerization of citrate to isocitrate via cis-aconitate | acnA | 4.2.1.3 | ko:K01681 |
| accessory/577/2/WP_132810337.1 | 391008.Smal_2353 | 0.0 | COG1629@1\|root,COG1629@2\|Bacteria,COG4771@2\|Bacteria,1MU9K@1224\|Proteobacteria,1RMTG@1236\|Gammaproteobacteria,1X4HX@135614\|Xanthomonadales | 135614\|Xanthomonadales | P | TonB-dependent receptor | - | - | - |
| accessory/588/2/WP_132810015.1 | 522373.Smlt2185 | 0.0 | COG3250@1\|root,COG3250@2\|Bacteria,1NYBH@1224\|Proteobacteria,1RZPC@1236\|Gammaproteobacteria,1XCMD@135614\|Xanthomonadales | 135614\|Xanthomonadales | G | Glycosyl hydrolases family 2 | - | 3.2.1.25 | ko:K01192 |
| accessory/595/2/WP_132810311.1 | 522373.Smlt2845 | 0.0 | COG4206@1\|root,COG4206@2\|Bacteria,1R47X@1224\|Proteobacteria,1T1X7@1236\|Gammaproteobacteria,1XD66@135614\|Xanthomonadales | 135614\|Xanthomonadales | H | TonB dependent receptor | - | - | ko:K02014 |
| accessory/664/2/WP_132810620.1 | 522373.Smlt3608 | 0.0 | COG1048@1\|root,COG1048@2\|Bacteria,1MU9T@1224\|Proteobacteria,1RN5I@1236\|Gammaproteobacteria,1X37M@135614\|Xanthomonadales | 135614\|Xanthomonadales | C | aconitate hydratase | acnA | 4.2.1.117 | ko:K20455 |
| accessory/676/2/WP_132809092.1 | 522373.Smlt0603 | 0.0 | COG2866@1\|root,COG2866@2\|Bacteria,1N9W9@1224\|Proteobacteria,1RPC7@1236\|Gammaproteobacteria,1X51W@135614\|Xanthomonadales | 135614\|Xanthomonadales | M | Zinc carboxypeptidase | - | - | - |
| accessory/696/2/WP_132809962.1 | 391008.Smal_1693 | 0.0 | COG1629@1\|root,COG4771@2\|Bacteria,1MUNK@1224\|Proteobacteria,1RN9S@1236\|Gammaproteobacteria,1X9QB@135614\|Xanthomonadales | 135614\|Xanthomonadales | M | TonB-dependent receptor | - | - | ko:K02014 |
| accessory/715/2/WP_132810450.1 | 391008.Smal_2577 | 0.0 | COG4993@1\|root,COG4993@2\|Bacteria,1MUQX@1224\|Proteobacteria,1RN5D@1236\|Gammaproteobacteria,1X42W@135614\|Xanthomonadales | 135614\|Xanthomonadales | G | Glucose dehydrogenase | gcd1 | 1.1.5.2 | ko:K00117 |
| accessory/754/2/WP_132809791.1 | 391008.Smal_1484 | 0.0 | COG1629@1\|root,COG4206@1\|root,COG1629@2\|Bacteria,COG4206@2\|Bacteria,1MXVP@1224\|Proteobacteria,1RS4C@1236\|Gammaproteobacteria,1X4VU@135614\|Xanthomonadales | 135614\|Xanthomonadales | HP | TonB-dependent receptor | - | - | - |
| accessory/781/2/WP_132810014.1 | 391008.Smal_1774 | 0.0 | COG3537@1\|root,COG3537@2\|Bacteria,1MXCY@1224\|Proteobacteria,1S0AN@1236\|Gammaproteobacteria,1X3FY@135614\|Xanthomonadales | 135614\|Xanthomonadales | G | Glycosyl hydrolase family 92 | - | - | - |
| accessory/787/2/WP_240792133.1 | 391008.Smal_2016 | 0.0 | COG4773@1\|root,COG4773@2\|Bacteria,1MW5E@1224\|Proteobacteria,1RMBD@1236\|Gammaproteobacteria,1X3Z3@135614\|Xanthomonadales | 135614\|Xanthomonadales | P | TonB dependent receptor | - | - | ko:K16088 |
| accessory/844/2/WP_132810957.1 | 522373.Smlt4410 | 0.0 | COG1629@1\|root,COG4771@2\|Bacteria,1MUNK@1224\|Proteobacteria,1RN9S@1236\|Gammaproteobacteria,1X4DH@135614\|Xanthomonadales | 135614\|Xanthomonadales | P | TonB-dependent receptor | - | - | ko:K02014 |
| accessory/847/2/WP_132810793.1 | 522373.Smlt4027 | 0.0 | COG3525@1\|root,COG3525@2\|Bacteria,1MVDE@1224\|Proteobacteria,1RMNI@1236\|Gammaproteobacteria,1X467@135614\|Xanthomonadales | 135614\|Xanthomonadales | G | Glycosyl hydrolase family 20, catalytic domain | nahA | 3.2.1.52 | ko:K12373 |
| accessory/873/2/WP_132809799.1 | 391008.Smal_1495 | 0.0 | COG4772@1\|root,COG4772@2\|Bacteria,1NTC4@1224\|Proteobacteria,1RPFQ@1236\|Gammaproteobacteria,1X4W7@135614\|Xanthomonadales | 135614\|Xanthomonadales | M | TonB-dependent receptor | - | - | - |
| accessory/893/2/WP_132809476.1 | 391008.Smal_1020 | 0.0 | COG3408@1\|root,COG3408@2\|Bacteria,1QWGV@1224\|Proteobacteria,1RYXV@1236\|Gammaproteobacteria,1X56X@135614\|Xanthomonadales | 135614\|Xanthomonadales | G | Glycogen debranching enzyme | - | - | - |
| accessory/904/2/WP_132810235.1 | 391008.Smal_2174 | 0.0 | COG3537@1\|root,COG3537@2\|Bacteria,1MXCY@1224\|Proteobacteria,1RYV7@1236\|Gammaproteobacteria,1XC59@135614\|Xanthomonadales | 135614\|Xanthomonadales | G | Hydrolase | - | - | - |
| accessory/921/2/WP_132808680.1 | 391008.Smal_0037 | 0.0 | COG1629@1\|root,COG4774@1\|root,COG1629@2\|Bacteria,COG4774@2\|Bacteria,1NMCN@1224\|Proteobacteria,1T2IM@1236\|Gammaproteobacteria,1XD6E@135614\|Xanthomonadales | 135614\|Xanthomonadales | P | TonB dependent receptor | - | - | ko:K02014 |
| accessory/953/2/WP_206138645.1 | 391008.Smal_2110 | 0.0 | COG4773@1\|root,COG4773@2\|Bacteria,1QTXJ@1224\|Proteobacteria,1T2K9@1236\|Gammaproteobacteria,1XD9B@135614\|Xanthomonadales | 135614\|Xanthomonadales | P | TonB dependent receptor | - | - | - |
| accessory/970/2/WP_132808932.1 | 391008.Smal_0331 | 0.0 | COG1472@1\|root,COG1472@2\|Bacteria,1MVIV@1224\|Proteobacteria,1RMA0@1236\|Gammaproteobacteria,1X4AX@135614\|Xanthomonadales | 135614\|Xanthomonadales | G | Belongs to the glycosyl hydrolase 3 family | bglX | 3.2.1.21 | ko:K05349 |
| accessory/1059/2/WP_132809538.1 | 391008.Smal_1082 | 0.0 | COG4773@1\|root,COG4773@2\|Bacteria,1MW5E@1224\|Proteobacteria,1RMBD@1236\|Gammaproteobacteria,1X9DR@135614\|Xanthomonadales | 135614\|Xanthomonadales | P | TonB dependent receptor | - | - | ko:K16088 |
| accessory/1065/2/WP_132809803.1 | 391008.Smal_1501 | 0.0 | COG4773@1\|root,COG4773@2\|Bacteria,1MW5E@1224\|Proteobacteria,1RMBD@1236\|Gammaproteobacteria,1X3NJ@135614\|Xanthomonadales | 135614\|Xanthomonadales | P | TonB-dependent siderophore receptor | - | - | ko:K16088 |
| accessory/1084/2/WP_132810225.1 | 391008.Smal_2161 | 0.0 | COG4773@1\|root,COG4773@2\|Bacteria,1QTXJ@1224\|Proteobacteria,1SM4W@1236\|Gammaproteobacteria | 1236\|Gammaproteobacteria | P | TonB dependent receptor | - | - | ko:K02014 |
| accessory/1102/2/WP_071229522.1 | 391008.Smal_3410 | 0.0 | COG4773@1\|root,COG4773@2\|Bacteria,1MW5E@1224\|Proteobacteria,1RMBD@1236\|Gammaproteobacteria,1X3NJ@135614\|Xanthomonadales | 135614\|Xanthomonadales | P | TonB-dependent siderophore receptor | fhuE | - | ko:K16088 |
| accessory/1109/2/WP_132809789.1 | 391008.Smal_1482 | 0.0 | COG4773@1\|root,COG4773@2\|Bacteria,1MW5E@1224\|Proteobacteria,1RMBD@1236\|Gammaproteobacteria,1X3NJ@135614\|Xanthomonadales | 1236\|Gammaproteobacteria | P | TonB-dependent siderophore receptor | - | - | ko:K16088 |
| accessory/1263/2/WP_132810279.1 | 1537715.JQFJ01000002_gene2533 | 5.85e-290 | COG1629@1\|root,COG4771@2\|Bacteria,1MUC1@1224\|Proteobacteria,2VFU5@28211\|Alphaproteobacteria,2KDZN@204457\|Sphingomonadales | 204457\|Sphingomonadales | P | TonB dependent receptor | - | - | - |
| accessory/1268/2/WP_049422603.1 | 391008.Smal_0797 | 0.0 | COG0446@1\|root,COG1902@1\|root,COG0446@2\|Bacteria,COG1902@2\|Bacteria,1MVE0@1224\|Proteobacteria,1RNM8@1236\|Gammaproteobacteria,1X445@135614\|Xanthomonadales | 135614\|Xanthomonadales | C | 2,4-dienoyl-coa reductase | fadH | 1.3.1.34 | ko:K00219 |
| accessory/1286/2/WP_049423773.1 | 522373.Smlt4135 | 0.0 | COG1629@1\|root,COG4771@2\|Bacteria,1MUC1@1224\|Proteobacteria,1RNHR@1236\|Gammaproteobacteria,1X5K3@135614\|Xanthomonadales | 135614\|Xanthomonadales | P | TonB-dependent receptor | - | - | ko:K16089 |
| accessory/5328/2/WP_049422074.1 | 325777.GW15_0207175 | 1.26e-194 | COG0667@1\|root,COG0667@2\|Bacteria,1PDY4@1224\|Proteobacteria,1RQYV@1236\|Gammaproteobacteria | 1236\|Gammaproteobacteria | C | Aldo Keto reductase | - | - | - |
| accessory/5331/2/WP_132810806.1 | 522373.Smlt4074 | 1.09e-228 | COG0673@1\|root,COG0673@2\|Bacteria,1MUZI@1224\|Proteobacteria,1RQV3@1236\|Gammaproteobacteria,1XCBH@135614\|Xanthomonadales | 135614\|Xanthomonadales | S | Oxidoreductase family, C-terminal alpha/beta domain | - | - | - |
| accessory/5334/2/WP_132809611.1 | 1500893.JQNB01000001_gene769 | 3.13e-107 | COG0438@1\|root,COG0438@2\|Bacteria,1N01H@1224\|Proteobacteria,1S8Z9@1236\|Gammaproteobacteria,1X5Z9@135614\|Xanthomonadales | 135614\|Xanthomonadales | M | glycosyltransferase | - | - | - |
| accessory/5337/2/WP_107431433.1 | 391008.Smal_0933 | 8.66e-229 | COG0667@1\|root,COG0667@2\|Bacteria,1MVEH@1224\|Proteobacteria,1RPX5@1236\|Gammaproteobacteria,1X56H@135614\|Xanthomonadales | 135614\|Xanthomonadales | C | oxidoreductases (related to aryl-alcohol dehydrogenases) | - | - | - |
| accessory/5359/2/WP_032127690.1 | 391008.Smal_2576 | 4.81e-227 | COG0604@1\|root,COG0604@2\|Bacteria,1MWBD@1224\|Proteobacteria,1RPCQ@1236\|Gammaproteobacteria,1X5U3@135614\|Xanthomonadales | 135614\|Xanthomonadales | C | Zinc-binding dehydrogenase | - | - | - |
| accessory/5375/2/WP_132811150.1 | 84531.JMTZ01000050_gene898 | 5.15e-92 | COG1216@1\|root,COG1216@2\|Bacteria,1QVRU@1224\|Proteobacteria,1T2IB@1236\|Gammaproteobacteria | 1236\|Gammaproteobacteria | S | Glycosyltransferase like family 2 | - | - | - |
| accessory/5418/2/WP_132809122.1 | 1123035.ARLA01000022_gene686 | 6.42e-94 | COG0463@1\|root,COG0463@2\|Bacteria,4NEVT@976\|Bacteroidetes,1HXT8@117743\|Flavobacteriia,4C375@83612\|Psychroflexus | 976\|Bacteroidetes | M | Glycosyl transferase family 2 | - | - | - |
| accessory/5433/2/WP_049447040.1 | 522373.Smlt4529 | 6.02e-220 | COG0667@1\|root,COG0667@2\|Bacteria,1MV2Y@1224\|Proteobacteria,1RQ1X@1236\|Gammaproteobacteria,1X4PU@135614\|Xanthomonadales | 135614\|Xanthomonadales | C | oxidoreductases (related to aryl-alcohol dehydrogenases) | - | - | - |
| accessory/5451/2/WP_049400603.1 | 522373.Smlt2843 | 3.77e-222 | COG0604@1\|root,COG0604@2\|Bacteria,1MXCV@1224\|Proteobacteria,1RSNU@1236\|Gammaproteobacteria,1X4ZY@135614\|Xanthomonadales | 135614\|Xanthomonadales | C | alcohol dehydrogenase | - | - | - |
| accessory/5476/2/WP_132809748.1 | 522373.Smlt1603 | 3.56e-206 | COG0741@1\|root,COG0741@2\|Bacteria,1MZ4X@1224\|Proteobacteria,1S8R3@1236\|Gammaproteobacteria,1XC67@135614\|Xanthomonadales | 135614\|Xanthomonadales | M | transglycosylase | - | - | - |
| accessory/5491/2/WP_132810648.1 | 743721.Psesu_0540 | 1.65e-107 | COG1216@1\|root,COG1216@2\|Bacteria,1QTEK@1224\|Proteobacteria,1S5EC@1236\|Gammaproteobacteria,1X6B6@135614\|Xanthomonadales | 135614\|Xanthomonadales | S | PFAM Glycosyl transferase family 2 | - | - | ko:K12990 |
| accessory/5539/2/WP_132810300.1 | 391008.Smal_2279 | 4.97e-220 | COG2141@1\|root,COG2141@2\|Bacteria,1MWDV@1224\|Proteobacteria,1RYCM@1236\|Gammaproteobacteria,1X967@135614\|Xanthomonadales | 135614\|Xanthomonadales | C | Luciferase-like monooxygenase | - | - | - |
| accessory/5559/2/WP_049400864.1 | 391008.Smal_3833 | 5.3e-215 | COG1171@1\|root,COG1171@2\|Bacteria,1MVWJ@1224\|Proteobacteria,1RPGU@1236\|Gammaproteobacteria,1X2XV@135614\|Xanthomonadales | 135614\|Xanthomonadales | E | dehydratase | ilvA | 4.3.1.19 | ko:K01754 |
| accessory/5901/2/WP_206138731.1 | 391008.Smal_3884 | 2.72e-165 | COG0451@1\|root,COG0451@2\|Bacteria,1MWVJ@1224\|Proteobacteria,1RNDT@1236\|Gammaproteobacteria,1X66F@135614\|Xanthomonadales | 135614\|Xanthomonadales | GM | NAD dependent epimerase/dehydratase family | - | - | - |
| accessory/6112/2/WP_132809496.1 | 391008.Smal_1041 | 1.53e-212 | COG3622@1\|root,COG3622@2\|Bacteria,1MV53@1224\|Proteobacteria,1RQF9@1236\|Gammaproteobacteria,1X4UB@135614\|Xanthomonadales | 135614\|Xanthomonadales | G | Xylose isomerase-like TIM barrel | - | 5.3.1.22 | ko:K01816 |
| accessory/6468/2/WP_132809168.1 | 391008.Smal_0557 | 1.4e-201 | COG1216@1\|root,COG1216@2\|Bacteria,1QP7Y@1224\|Proteobacteria,1RR12@1236\|Gammaproteobacteria,1X4AI@135614\|Xanthomonadales | 135614\|Xanthomonadales | S | glycosyl transferase family 2 | - | - | - |
| accessory/6478/2/WP_132809501.1 | 391008.Smal_1046 | 1.2e-183 | COG1082@1\|root,COG1082@2\|Bacteria,1RAC3@1224\|Proteobacteria,1RS39@1236\|Gammaproteobacteria,1X6BI@135614\|Xanthomonadales | 135614\|Xanthomonadales | G | xylose isomerase | - | - | - |
| accessory/6551/2/WP_132810658.1 | 522373.Smlt3697 | 8.6e-182 | COG2819@1\|root,COG2819@2\|Bacteria,1R9SG@1224\|Proteobacteria,1T1D1@1236\|Gammaproteobacteria,1X6IU@135614\|Xanthomonadales | 135614\|Xanthomonadales | S | Putative esterase | - | - | ko:K07017 |
| accessory/6663/2/WP_071228917.1 | 391008.Smal_1727 | 1.85e-169 | COG1028@1\|root,COG1028@2\|Bacteria,1MW46@1224\|Proteobacteria,1S4NN@1236\|Gammaproteobacteria,1XCB8@135614\|Xanthomonadales | 135614\|Xanthomonadales | IQ | short chain dehydrogenase | - | 1.1.1.100 | ko:K00059 |
| accessory/6874/2/WP_071229432.1 | 391008.Smal_3222 | 7.76e-190 | COG0491@1\|root,COG0491@2\|Bacteria,1MURA@1224\|Proteobacteria,1RN27@1236\|Gammaproteobacteria,1X3E4@135614\|Xanthomonadales | 135614\|Xanthomonadales | S | Zn-dependent hydrolases including glyoxylases | - | - | - |
| accessory/6880/2/WP_049424806.1 | 391008.Smal_0432 | 4.35e-194 | COG3384@1\|root,COG3384@2\|Bacteria,1MXJZ@1224\|Proteobacteria,1RR5P@1236\|Gammaproteobacteria,1X3A0@135614\|Xanthomonadales | 135614\|Xanthomonadales | S | dioxygenase | - | - | ko:K15777 |
| accessory/7013/2/WP_132810809.1 | 522373.Smlt4092 | 9.25e-178 | COG1028@1\|root,COG1028@2\|Bacteria,1MUAY@1224\|Proteobacteria,1S1XJ@1236\|Gammaproteobacteria,1X4DQ@135614\|Xanthomonadales | 135614\|Xanthomonadales | IQ | short-chain dehydrogenase reductase | - | - | - |
| accessory/7042/2/WP_019336772.1 | 522373.Smlt4539 | 6.07e-184 | COG0656@1\|root,COG0656@2\|Bacteria,1MWFS@1224\|Proteobacteria,1RMX6@1236\|Gammaproteobacteria,1X6T4@135614\|Xanthomonadales | 135614\|Xanthomonadales | S | Catalyzes the reduction of 2,5-diketo-D-gluconic acid to 2-keto-L-gulonic acid | dkgB | 1.1.1.346 | ko:K06222 |
| accessory/7139/2/WP_132808833.1 | 522373.Smlt0266 | 1.05e-182 | COG1024@1\|root,COG1024@2\|Bacteria,1MWZC@1224\|Proteobacteria,1RR3Z@1236\|Gammaproteobacteria,1X4HN@135614\|Xanthomonadales | 135614\|Xanthomonadales | I | Belongs to the enoyl-CoA hydratase isomerase family | paaF | - | - |
| accessory/7147/2/WP_049461643.1 | 391008.Smal_3694 | 3.34e-174 | COG1028@1\|root,COG1028@2\|Bacteria,1MUWC@1224\|Proteobacteria,1RQBN@1236\|Gammaproteobacteria,1X4C0@135614\|Xanthomonadales | 135614\|Xanthomonadales | IQ | Belongs to the short-chain dehydrogenases reductases (SDR) family | - | - | - |
| accessory/7154/2/WP_049399918.1 | 391008.Smal_1375 | 1.1e-160 | COG1028@1\|root,COG1028@2\|Bacteria,1MWBC@1224\|Proteobacteria,1RNNV@1236\|Gammaproteobacteria,1X3SZ@135614\|Xanthomonadales | 135614\|Xanthomonadales | IQ | short-chain dehydrogenase | - | - | - |
| accessory/7318/2/WP_132810977.1 | 391008.Smal_3830 | 6.8e-176 | COG1028@1\|root,COG1028@2\|Bacteria,1MWJI@1224\|Proteobacteria,1RY5G@1236\|Gammaproteobacteria,1X7AZ@135614\|Xanthomonadales | 135614\|Xanthomonadales | IQ | Enoyl-(Acyl carrier protein) reductase | - | 1.1.1.100 | ko:K00059 |
| accessory/7322/2/WP_132809956.1 | 391008.Smal_1682 | 6.46e-170 | COG1028@1\|root,COG1028@2\|Bacteria,1MU5Y@1224\|Proteobacteria,1RP7D@1236\|Gammaproteobacteria,1X464@135614\|Xanthomonadales | 135614\|Xanthomonadales | IQ | Belongs to the short-chain dehydrogenases reductases (SDR) family | hbdH1 | 1.1.1.30 | ko:K00019 |
| accessory/7358/2/WP_132811131.1 | 391008.Smal_1809 | 1.6e-174 | COG1028@1\|root,COG1028@2\|Bacteria,1ND2U@1224\|Proteobacteria,1RZZH@1236\|Gammaproteobacteria,1X5HF@135614\|Xanthomonadales | 135614\|Xanthomonadales | IQ | dehydrogenase reductase | - | 1.1.1.100 | ko:K00059 |
| accessory/7412/2/WP_132811023.1 | 391008.Smal_3919 | 1.63e-186 | COG1216@1\|root,COG1216@2\|Bacteria,1QVEM@1224\|Proteobacteria,1T2CN@1236\|Gammaproteobacteria,1X4G9@135614\|Xanthomonadales | 135614\|Xanthomonadales | S | Glycosyl transferase family 2 | - | - | - |
| accessory/1163/2/WP_165929924.1 | 391008.Smal_0907 | 0.0 | COG4774@1\|root,COG4774@2\|Bacteria,1MV0X@1224\|Proteobacteria,1S1IP@1236\|Gammaproteobacteria,1XCG9@135614\|Xanthomonadales | 135614\|Xanthomonadales | M | TonB dependent receptor | - | - | ko:K02014 |
| accessory/1176/2/WP_240792137.1 | 1458357.BG58_03305 | 2.71e-193 | COG0438@1\|root,COG1215@1\|root,COG0438@2\|Bacteria,COG1215@2\|Bacteria,1RD9X@1224\|Proteobacteria,2WDD3@28216\|Betaproteobacteria,1KA1R@119060\|Burkholderiaceae | 28216\|Betaproteobacteria | M | Glycosyl transferases group 1 | - | - | - |
| accessory/1191/2/WP_132810435.1 | 522373.Smlt3115 | 0.0 | COG4772@1\|root,COG4772@2\|Bacteria,1MUIH@1224\|Proteobacteria,1RZ1Q@1236\|Gammaproteobacteria,1X5KK@135614\|Xanthomonadales | 135614\|Xanthomonadales | P | TonB-dependent receptor | - | - | ko:K02014 |
| accessory/1263/2/WP_132810279.1 | 1537715.JQFJ01000002_gene2533 | 5.85e-290 | COG1629@1\|root,COG4771@2\|Bacteria,1MUC1@1224\|Proteobacteria,2VFU5@28211\|Alphaproteobacteria,2KDZN@204457\|Sphingomonadales | 204457\|Sphingomonadales | P | TonB dependent receptor | - | - | - |
| accessory/1268/2/WP_049422603.1 | 391008.Smal_0797 | 0.0 | COG0446@1\|root,COG1902@1\|root,COG0446@2\|Bacteria,COG1902@2\|Bacteria,1MVE0@1224\|Proteobacteria,1RNM8@1236\|Gammaproteobacteria,1X445@135614\|Xanthomonadales | 135614\|Xanthomonadales | C | 2,4-dienoyl-coa reductase | fadH | 1.3.1.34 | ko:K00219 |
| accessory/1286/2/WP_049423773.1 | 522373.Smlt4135 | 0.0 | COG1629@1\|root,COG4771@2\|Bacteria,1MUC1@1224\|Proteobacteria,1RNHR@1236\|Gammaproteobacteria,1X5K3@135614\|Xanthomonadales | 135614\|Xanthomonadales | P | TonB-dependent receptor | - | - | ko:K16089 |
| accessory/1726/2/WP_132810673.1 | 59538.XP_005977212.1 | 0.0 | COG3387@1\|root,2QS7A@2759\|Eukaryota,38D5C@33154\|Opisthokonta | 33154\|Opisthokonta | G | Glycosyl hydrolases family 15 | - | - | - |
| accessory/2012/2/WP_132810589.1 | 522373.Smlt3528 | 0.0 | COG1960@1\|root,COG1960@2\|Bacteria,1MU20@1224\|Proteobacteria,1RN7X@1236\|Gammaproteobacteria,1X3MF@135614\|Xanthomonadales | 135614\|Xanthomonadales | I | acyl-coa dehydrogenase | - | - | ko:K09456 |
| accessory/2041/2/WP_132809297.1 | 522373.Smlt0940 | 0.0 | COG0154@1\|root,COG0154@2\|Bacteria,1MUVQ@1224\|Proteobacteria,1RP7E@1236\|Gammaproteobacteria,1X4K6@135614\|Xanthomonadales | 135614\|Xanthomonadales | J | Catalyzes the hydrolysis of a monocarboxylic acid amid to form a monocarboxylate and ammonia | gatAX | 3.5.1.4 | ko:K01426 |
| accessory/2114/2/WP_132810561.1 | 391008.Smal_2878 | 0.0 | COG1233@1\|root,COG1233@2\|Bacteria,1MV2R@1224\|Proteobacteria,1RSGI@1236\|Gammaproteobacteria,1X5MS@135614\|Xanthomonadales | 135614\|Xanthomonadales | Q | Oxidoreductase | - | - | - |
| accessory/2270/2/WP_239509194.1 | 522373.Smlt1793 | 0.0 | COG0364@1\|root,COG0364@2\|Bacteria,1MUN0@1224\|Proteobacteria,1RN76@1236\|Gammaproteobacteria,1X49Y@135614\|Xanthomonadales | 135614\|Xanthomonadales | G | Catalyzes the oxidation of glucose 6-phosphate to 6- phosphogluconolactone | zwf | 1.1.1.363,1.1.1.49 | ko:K00036 |
| accessory/2477/2/WP_132810292.1 | 391008.Smal_2270 | 0.0 | COG4222@1\|root,COG4222@2\|Bacteria,1MVDD@1224\|Proteobacteria,1RYXE@1236\|Gammaproteobacteria,1X7TW@135614\|Xanthomonadales | 135614\|Xanthomonadales | S | Esterase-like activity of phytase | - | - | - |
| accessory/2836/2/WP_132809189.1 | 391008.Smal_0587 | 0.0 | COG0277@1\|root,COG0277@2\|Bacteria,1MU6Y@1224\|Proteobacteria,1RQX2@1236\|Gammaproteobacteria,1X3SQ@135614\|Xanthomonadales | 135614\|Xanthomonadales | C | FAD FMN-containing dehydrogenases | dld | 1.1.2.4,1.1.5.12 | ko:K00102,ko:K03777 |
| accessory/3065/2/WP_071228936.1 | 391008.Smal_1697 | 0.0 | COG2141@1\|root,COG2141@2\|Bacteria,1MUVN@1224\|Proteobacteria,1RQ78@1236\|Gammaproteobacteria,1X5Q3@135614\|Xanthomonadales | 135614\|Xanthomonadales | C | Nitrilotriacetate monooxygenase | - | - | - |
| accessory/3072/2/WP_132810814.1 | 522373.Smlt4105 | 1.09e-306 | COG2204@1\|root,COG2204@2\|Bacteria,1MU0N@1224\|Proteobacteria,1RMCK@1236\|Gammaproteobacteria,1X3II@135614\|Xanthomonadales | 135614\|Xanthomonadales | T | CheY-like receiver AAA-type ATPase and DNA-binding domains | - | - | - |
| accessory/3441/2/WP_132811020.1 | 391008.Smal_3913 | 8.51e-306 | COG0644@1\|root,COG0644@2\|Bacteria,1MZVI@1224\|Proteobacteria,1RMNS@1236\|Gammaproteobacteria,1X4V2@135614\|Xanthomonadales | 135614\|Xanthomonadales | C | COG0644 Dehydrogenases (flavoproteins) | - | - | - |
| accessory/3580/2/WP_032128677.1 | 391008.Smal_0042 | 2.95e-282 | COG1398@1\|root,COG1398@2\|Bacteria,1N2MA@1224\|Proteobacteria,1RSN1@1236\|Gammaproteobacteria,1X377@135614\|Xanthomonadales | 135614\|Xanthomonadales | I | desaturase | - | 1.14.19.1 | ko:K00507 |
| accessory/3992/2/WP_071229412.1 | 522373.Smlt0265 | 3.96e-274 | COG1960@1\|root,COG1960@2\|Bacteria,1MUDR@1224\|Proteobacteria,1RMMJ@1236\|Gammaproteobacteria,1X56A@135614\|Xanthomonadales | 135614\|Xanthomonadales | I | Acyl-CoA dehydrogenase | - | - | - |
| accessory/4001/2/WP_132808835.1 | 391008.Smal_0222 | 1.34e-278 | COG1024@1\|root,COG1024@2\|Bacteria,1MU0B@1224\|Proteobacteria,1RN07@1236\|Gammaproteobacteria,1X5E3@135614\|Xanthomonadales | 135614\|Xanthomonadales | I | Enoyl-CoA hydratase | - | - | - |
| accessory/4023/2/WP_132808935.1 | 522373.Smlt0451 | 1.15e-278 | COG2133@1\|root,COG2133@2\|Bacteria,1MV2E@1224\|Proteobacteria,1RNGN@1236\|Gammaproteobacteria,1X362@135614\|Xanthomonadales | 135614\|Xanthomonadales | G | Dehydrogenase | yliI | - | - |
| accessory/4079/2/WP_024957648.1 | 522373.Smlt2831 | 1.72e-290 | COG3239@1\|root,COG3239@2\|Bacteria,1QKDT@1224\|Proteobacteria,1RQCX@1236\|Gammaproteobacteria,1X49C@135614\|Xanthomonadales | 135614\|Xanthomonadales | I | desaturase | - | 1.14.19.3 | ko:K00508 |
| accessory/4231/2/WP_032129074.1 | 391008.Smal_0818 | 3.34e-267 | COG0438@1\|root,COG0438@2\|Bacteria,1MUB7@1224\|Proteobacteria,1RQYE@1236\|Gammaproteobacteria,1X41B@135614\|Xanthomonadales | 135614\|Xanthomonadales | M | Glycosyl transferase | - | - | - |
| accessory/4297/2/WP_125892196.1 | 391008.Smal_3580 | 4.83e-240 | COG0438@1\|root,COG0438@2\|Bacteria,1Q8II@1224\|Proteobacteria,1S5T1@1236\|Gammaproteobacteria,1X9W5@135614\|Xanthomonadales | 135614\|Xanthomonadales | M | Glycosyltransferase Family 4 | - | - | - |
| accessory/4487/2/WP_132808821.1 | 522373.Smlt0253 | 1.72e-266 | COG1902@1\|root,COG1902@2\|Bacteria,1MVE0@1224\|Proteobacteria,1RMII@1236\|Gammaproteobacteria,1X3AF@135614\|Xanthomonadales | 135614\|Xanthomonadales | C | Oxidoreductase | - | - | - |
| accessory/4490/2/WP_165929936.1 | 522373.Smlt2832 | 3.62e-257 | COG1018@1\|root,COG1018@2\|Bacteria,1MY2Q@1224\|Proteobacteria,1S01V@1236\|Gammaproteobacteria,1X4DC@135614\|Xanthomonadales | 135614\|Xanthomonadales | C | Oxidoreductase | - | - | - |
| accessory/4749/2/WP_032129891.1 | 522373.Smlt3987 | 7.82e-265 | COG1064@1\|root,COG1064@2\|Bacteria,1MUTT@1224\|Proteobacteria,1RN4D@1236\|Gammaproteobacteria,1X3TU@135614\|Xanthomonadales | 135614\|Xanthomonadales | S | dehydrogenase | - | - | ko:K13979 |
| accessory/4913/2/WP_132810797.1 | 391008.Smal_3437 | 4.94e-245 | COG0673@1\|root,COG0673@2\|Bacteria,1MU8F@1224\|Proteobacteria,1RNKY@1236\|Gammaproteobacteria,1X4MJ@135614\|Xanthomonadales | 135614\|Xanthomonadales | S | Oxidoreductase | ydgJ | - | - |
| accessory/5024/2/WP_132808672.1 | 391008.Smal_0028 | 3.64e-249 | COG1064@1\|root,COG1064@2\|Bacteria,1MUTT@1224\|Proteobacteria,1RPQC@1236\|Gammaproteobacteria,1X4TA@135614\|Xanthomonadales | 135614\|Xanthomonadales | S | alcohol dehydrogenase | - | 1.1.1.1 | ko:K13953 |
| accessory/5291/2/WP_132808866.1 | 522373.Smlt0347 | 1.05e-225 | COG0491@1\|root,COG0491@2\|Bacteria,1MXKX@1224\|Proteobacteria,1RR31@1236\|Gammaproteobacteria,1X47R@135614\|Xanthomonadales | 135614\|Xanthomonadales | S | Zn-dependent hydrolases including glyoxylases | - | - | - |
| accessory/5317/2/WP_132809964.1 | 391008.Smal_1696 | 2.89e-226 | COG2141@1\|root,COG2141@2\|Bacteria,1MVF0@1224\|Proteobacteria,1RPK5@1236\|Gammaproteobacteria,1X8VK@135614\|Xanthomonadales | 135614\|Xanthomonadales | C | Luciferase-like monooxygenase | - | - | - |
| accessory/5328/2/WP_049422074.1 | 325777.GW15_0207175 | 1.26e-194 | COG0667@1\|root,COG0667@2\|Bacteria,1PDY4@1224\|Proteobacteria,1RQYV@1236\|Gammaproteobacteria | 1236\|Gammaproteobacteria | C | Aldo Keto reductase | - | - | - |
| accessory/5334/2/WP_132809611.1 | 1500893.JQNB01000001_gene769 | 3.13e-107 | COG0438@1\|root,COG0438@2\|Bacteria,1N01H@1224\|Proteobacteria,1S8Z9@1236\|Gammaproteobacteria,1X5Z9@135614\|Xanthomonadales | 135614\|Xanthomonadales | M | glycosyltransferase | - | - | - |
| accessory/5337/2/WP_107431433.1 | 391008.Smal_0933 | 8.66e-229 | COG0667@1\|root,COG0667@2\|Bacteria,1MVEH@1224\|Proteobacteria,1RPX5@1236\|Gammaproteobacteria,1X56H@135614\|Xanthomonadales | 135614\|Xanthomonadales | C | oxidoreductases (related to aryl-alcohol dehydrogenases) | - | - | - |
| accessory/5359/2/WP_032127690.1 | 391008.Smal_2576 | 4.81e-227 | COG0604@1\|root,COG0604@2\|Bacteria,1MWBD@1224\|Proteobacteria,1RPCQ@1236\|Gammaproteobacteria,1X5U3@135614\|Xanthomonadales | 135614\|Xanthomonadales | C | Zinc-binding dehydrogenase | - | - | - |
| accessory/5375/2/WP_132811150.1 | 84531.JMTZ01000050_gene898 | 5.15e-92 | COG1216@1\|root,COG1216@2\|Bacteria,1QVRU@1224\|Proteobacteria,1T2IB@1236\|Gammaproteobacteria | 1236\|Gammaproteobacteria | S | Glycosyltransferase like family 2 | - | - | - |
| accessory/5418/2/WP_132809122.1 | 1123035.ARLA01000022_gene686 | 6.42e-94 | COG0463@1\|root,COG0463@2\|Bacteria,4NEVT@976\|Bacteroidetes,1HXT8@117743\|Flavobacteriia,4C375@83612\|Psychroflexus | 976\|Bacteroidetes | M | Glycosyl transferase family 2 | - | - | - |
| accessory/5451/2/WP_049400603.1 | 522373.Smlt2843 | 3.77e-222 | COG0604@1\|root,COG0604@2\|Bacteria,1MXCV@1224\|Proteobacteria,1RSNU@1236\|Gammaproteobacteria,1X4ZY@135614\|Xanthomonadales | 135614\|Xanthomonadales | C | alcohol dehydrogenase | - | - | - |
| accessory/5476/2/WP_132809748.1 | 522373.Smlt1603 | 3.56e-206 | COG0741@1\|root,COG0741@2\|Bacteria,1MZ4X@1224\|Proteobacteria,1S8R3@1236\|Gammaproteobacteria,1XC67@135614\|Xanthomonadales | 135614\|Xanthomonadales | M | transglycosylase | - | - | - |
| accessory/5539/2/WP_132810300.1 | 391008.Smal_2279 | 4.97e-220 | COG2141@1\|root,COG2141@2\|Bacteria,1MWDV@1224\|Proteobacteria,1RYCM@1236\|Gammaproteobacteria,1X967@135614\|Xanthomonadales | 135614\|Xanthomonadales | C | Luciferase-like monooxygenase | - | - | - |
| accessory/5547/2/WP_132810097.1 | 522373.Smlt2374 | 1.49e-203 | COG1398@1\|root,COG1398@2\|Bacteria,1N2MA@1224\|Proteobacteria,1RM88@1236\|Gammaproteobacteria,1X4PF@135614\|Xanthomonadales | 135614\|Xanthomonadales | I | fatty acid desaturase | desC | 1.14.19.1 | ko:K00507 |
| accessory/5559/2/WP_049400864.1 | 391008.Smal_3833 | 5.3e-215 | COG1171@1\|root,COG1171@2\|Bacteria,1MVWJ@1224\|Proteobacteria,1RPGU@1236\|Gammaproteobacteria,1X2XV@135614\|Xanthomonadales | 135614\|Xanthomonadales | E | dehydratase | ilvA | 4.3.1.19 | ko:K01754 |
| accessory/5659/2/WP_132810127.1 | 391008.Smal_2004 | 3.98e-229 | COG2267@1\|root,COG2267@2\|Bacteria,1MUG8@1224\|Proteobacteria,1RMFE@1236\|Gammaproteobacteria,1X43J@135614\|Xanthomonadales | 135614\|Xanthomonadales | I | Alpha beta hydrolase | - | 1.11.1.10 | ko:K00433 |
| accessory/5736/2/WP_132810657.1 | 391008.Smal_3111 | 2.1e-217 | COG2819@1\|root,COG2819@2\|Bacteria,1RB1X@1224\|Proteobacteria,1RY80@1236\|Gammaproteobacteria,1XC1G@135614\|Xanthomonadales | 135614\|Xanthomonadales | S | Putative esterase | - | - | ko:K07017 |
| accessory/5806/2/WP_132809384.1 | 391008.Smal_0915 | 2.54e-206 | COG0583@1\|root,COG0583@2\|Bacteria,1MW16@1224\|Proteobacteria,1RS6U@1236\|Gammaproteobacteria,1X5HU@135614\|Xanthomonadales | 135614\|Xanthomonadales | K | LysR family | - | - | - |
| accessory/5819/2/WP_107432530.1 | 522373.Smlt0522 | 8.03e-214 | COG0583@1\|root,COG0583@2\|Bacteria,1MW16@1224\|Proteobacteria,1RS6U@1236\|Gammaproteobacteria,1X5HU@135614\|Xanthomonadales | 135614\|Xanthomonadales | K | LysR family | - | - | - |
| accessory/5824/2/WP_132809207.1 | 391008.Smal_0615 | 7.78e-191 | COG0596@1\|root,COG0596@2\|Bacteria,1P1S6@1224\|Proteobacteria,1T49K@1236\|Gammaproteobacteria,1X9FI@135614\|Xanthomonadales | 135614\|Xanthomonadales | S | Alpha beta hydrolase | - | - | - |
| accessory/5874/2/WP_014648435.1 | 522373.Smlt4042 | 6.74e-213 | COG1023@1\|root,COG1023@2\|Bacteria,1QU14@1224\|Proteobacteria,1T1KM@1236\|Gammaproteobacteria,1X4EG@135614\|Xanthomonadales | 135614\|Xanthomonadales | G | 6-phosphogluconate dehydrogenase | gnd | 1.1.1.343,1.1.1.44 | ko:K00033 |
| accessory/5901/2/WP_206138731.1 | 391008.Smal_3884 | 2.72e-165 | COG0451@1\|root,COG0451@2\|Bacteria,1MWVJ@1224\|Proteobacteria,1RNDT@1236\|Gammaproteobacteria,1X66F@135614\|Xanthomonadales | 135614\|Xanthomonadales | GM | NAD dependent epimerase/dehydratase family | - | - | - |
| accessory/6150/2/WP_132811129.1 | 522373.Smlt2114 | 1.02e-202 | COG0583@1\|root,COG0583@2\|Bacteria,1MW16@1224\|Proteobacteria,1RS6U@1236\|Gammaproteobacteria,1X5HU@135614\|Xanthomonadales | 135614\|Xanthomonadales | K | LysR family | - | - | - |
| accessory/6422/2/WP_049401006.1 | 522373.Smlt4172 | 6.38e-192 | COG0463@1\|root,COG0463@2\|Bacteria,1PVP4@1224\|Proteobacteria,1RQUH@1236\|Gammaproteobacteria,1XD5Y@135614\|Xanthomonadales | 135614\|Xanthomonadales | M | Glycosyl transferase family 2 | - | - | - |
| accessory/6467/2/WP_132810734.1 | 522373.Smlt3879 | 4.13e-188 | COG1028@1\|root,COG1028@2\|Bacteria,1MUSQ@1224\|Proteobacteria,1RQQ9@1236\|Gammaproteobacteria,1X32T@135614\|Xanthomonadales | 135614\|Xanthomonadales | IQ | dehydrogenase | - | - | ko:K13775 |
| accessory/6468/2/WP_132809168.1 | 391008.Smal_0557 | 1.4e-201 | COG1216@1\|root,COG1216@2\|Bacteria,1QP7Y@1224\|Proteobacteria,1RR12@1236\|Gammaproteobacteria,1X4AI@135614\|Xanthomonadales | 135614\|Xanthomonadales | S | glycosyl transferase family 2 | - | - | - |
| accessory/6478/2/WP_132809501.1 | 391008.Smal_1046 | 1.2e-183 | COG1082@1\|root,COG1082@2\|Bacteria,1RAC3@1224\|Proteobacteria,1RS39@1236\|Gammaproteobacteria,1X6BI@135614\|Xanthomonadales | 135614\|Xanthomonadales | G | xylose isomerase | - | - | - |
| accessory/6874/2/WP_071229432.1 | 391008.Smal_3222 | 7.76e-190 | COG0491@1\|root,COG0491@2\|Bacteria,1MURA@1224\|Proteobacteria,1RN27@1236\|Gammaproteobacteria,1X3E4@135614\|Xanthomonadales | 135614\|Xanthomonadales | S | Zn-dependent hydrolases including glyoxylases | - | - | - |
| accessory/6880/2/WP_049424806.1 | 391008.Smal_0432 | 4.35e-194 | COG3384@1\|root,COG3384@2\|Bacteria,1MXJZ@1224\|Proteobacteria,1RR5P@1236\|Gammaproteobacteria,1X3A0@135614\|Xanthomonadales | 135614\|Xanthomonadales | S | dioxygenase | - | - | ko:K15777 |
| accessory/7013/2/WP_132810809.1 | 522373.Smlt4092 | 9.25e-178 | COG1028@1\|root,COG1028@2\|Bacteria,1MUAY@1224\|Proteobacteria,1S1XJ@1236\|Gammaproteobacteria,1X4DQ@135614\|Xanthomonadales | 135614\|Xanthomonadales | IQ | short-chain dehydrogenase reductase | - | - | - |
| accessory/7195/2/WP_132810093.1 | 522373.Smlt2369 | 3.01e-180 | COG3752@1\|root,COG3752@2\|Bacteria,1MXCP@1224\|Proteobacteria,1S26M@1236\|Gammaproteobacteria,1XBWB@135614\|Xanthomonadales | 135614\|Xanthomonadales | S | 3-oxo-5-alpha-steroid 4-dehydrogenase | - | - | - |
| accessory/7318/2/WP_132810977.1 | 391008.Smal_3830 | 6.8e-176 | COG1028@1\|root,COG1028@2\|Bacteria,1MWJI@1224\|Proteobacteria,1RY5G@1236\|Gammaproteobacteria,1X7AZ@135614\|Xanthomonadales | 135614\|Xanthomonadales | IQ | Enoyl-(Acyl carrier protein) reductase | - | 1.1.1.100 | ko:K00059 |
| accessory/7322/2/WP_132809956.1 | 391008.Smal_1682 | 6.46e-170 | COG1028@1\|root,COG1028@2\|Bacteria,1MU5Y@1224\|Proteobacteria,1RP7D@1236\|Gammaproteobacteria,1X464@135614\|Xanthomonadales | 135614\|Xanthomonadales | IQ | Belongs to the short-chain dehydrogenases reductases (SDR) family | hbdH1 | 1.1.1.30 | ko:K00019 |
| accessory/7358/2/WP_132811131.1 | 391008.Smal_1809 | 1.6e-174 | COG1028@1\|root,COG1028@2\|Bacteria,1ND2U@1224\|Proteobacteria,1RZZH@1236\|Gammaproteobacteria,1X5HF@135614\|Xanthomonadales | 135614\|Xanthomonadales | IQ | dehydrogenase reductase | - | 1.1.1.100 | ko:K00059 |
| accessory/7388/2/WP_024958208.1 | 391008.Smal_3879 | 1.24e-182 | COG1018@1\|root,COG1018@2\|Bacteria,1REP4@1224\|Proteobacteria,1S7ZN@1236\|Gammaproteobacteria,1XCEZ@135614\|Xanthomonadales | 135614\|Xanthomonadales | C | Oxidoreductase FAD-binding domain | - | - | - |
| accessory/7412/2/WP_132811023.1 | 391008.Smal_3919 | 1.63e-186 | COG1216@1\|root,COG1216@2\|Bacteria,1QVEM@1224\|Proteobacteria,1T2CN@1236\|Gammaproteobacteria,1X4G9@135614\|Xanthomonadales | 135614\|Xanthomonadales | S | Glycosyl transferase family 2 | - | - | - |
| accessory/7439/2/WP_049423570.1 | 522373.Smlt1829 | 2.91e-180 | COG1028@1\|root,COG1028@2\|Bacteria,1NHJ3@1224\|Proteobacteria,1SUFK@1236\|Gammaproteobacteria,1X6T1@135614\|Xanthomonadales | 135614\|Xanthomonadales | IQ | short chain dehydrogenase | - | - | - |
| accessory/7532/2/WP_132810294.1 | 391008.Smal_2271 | 3.59e-163 | COG1028@1\|root,COG1028@2\|Bacteria,1MUD2@1224\|Proteobacteria,1RN5N@1236\|Gammaproteobacteria,1XA2G@135614\|Xanthomonadales | 135614\|Xanthomonadales | IQ | NAD dependent epimerase/dehydratase family | - | 1.3.1.28 | ko:K00216 |
| accessory/7598/2/WP_049400632.1 | 391008.Smal_3069 | 1.9e-158 | COG1028@1\|root,COG1028@2\|Bacteria,1QWBB@1224\|Proteobacteria,1S1SF@1236\|Gammaproteobacteria,1XCBG@135614\|Xanthomonadales | 135614\|Xanthomonadales | IQ | NAD dependent epimerase/dehydratase family | - | 1.1.1.100 | ko:K00059 |
| accessory/7668/2/WP_132810231.1 | 391008.Smal_2168 | 5.06e-156 | COG1028@1\|root,COG1028@2\|Bacteria,1MU73@1224\|Proteobacteria,1RR5Q@1236\|Gammaproteobacteria,1X44T@135614\|Xanthomonadales | 135614\|Xanthomonadales | IQ | NAD dependent epimerase/dehydratase family | - | 1.1.1.100 | ko:K00059 |
| accessory/7702/2/WP_107433349.1 | 391008.Smal_1292 | 5.23e-154 | COG0730@1\|root,COG0730@2\|Bacteria,1MWAN@1224\|Proteobacteria,1S158@1236\|Gammaproteobacteria,1XBZ0@135614\|Xanthomonadales | 135614\|Xanthomonadales | S | Sulfite exporter TauE/SafE | - | - | ko:K07090 |
| accessory/7921/2/WP_132811014.1 | 522373.Smlt4544 | 1.74e-156 | COG2021@1\|root,COG2021@2\|Bacteria,1RDEH@1224\|Proteobacteria,1S75E@1236\|Gammaproteobacteria,1X5Y9@135614\|Xanthomonadales | 135614\|Xanthomonadales | E | Alpha beta hydrolase | - | - | - |
| accessory/7925/2/WP_032129364.1 | 391008.Smal_2972 | 1.74e-165 | COG0745@1\|root,COG0745@2\|Bacteria,1RHTT@1224\|Proteobacteria,1T1MZ@1236\|Gammaproteobacteria,1X6CU@135614\|Xanthomonadales | 135614\|Xanthomonadales | T | COG0784 FOG CheY-like receiver | - | - | - |
| accessory/8056/2/WP_165929920.1 | 1366050.N234_16615 | 1.29e-53 | COG0463@1\|root,COG0463@2\|Bacteria,1N4WK@1224\|Proteobacteria,2WHRC@28216\|Betaproteobacteria,1KDHA@119060\|Burkholderiaceae | 28216\|Betaproteobacteria | M | Glycosyl transferase family 2 | - | - | - |
| accessory/8106/2/WP_071228168.1 | 391008.Smal_3308 | 3.35e-157 | COG0810@1\|root,COG0810@2\|Bacteria,1MZXC@1224\|Proteobacteria,1SA2U@1236\|Gammaproteobacteria,1X8JV@135614\|Xanthomonadales | 135614\|Xanthomonadales | M | TonB C terminal | - | - | ko:K03832 |
| accessory/8115/2/WP_132809972.1 | 522373.Smlt2116 | 1.26e-156 | COG1028@1\|root,COG1028@2\|Bacteria,1QW9C@1224\|Proteobacteria,1S0NM@1236\|Gammaproteobacteria | 1236\|Gammaproteobacteria | IQ | dehydrogenase reductase | - | - | - |
| accessory/8147/2/WP_049398426.1 | 522373.Smlt2939 | 3.49e-154 | COG0810@1\|root,COG0810@2\|Bacteria,1PFM3@1224\|Proteobacteria,1SWJ2@1236\|Gammaproteobacteria,1X8UH@135614\|Xanthomonadales | 135614\|Xanthomonadales | M | TonB C terminal | - | - | ko:K03832 |
| accessory/8180/2/WP_132810668.1 | 522373.Smlt3734 | 5e-162 | COG0625@1\|root,COG0625@2\|Bacteria,1N8XH@1224\|Proteobacteria,1SJPR@1236\|Gammaproteobacteria,1X3U2@135614\|Xanthomonadales | 135614\|Xanthomonadales | O | Glutathione S-transferase | - | 2.5.1.18 | ko:K00799 |
| accessory/8182/2/WP_132811116.1 | 522373.Smlt1229 | 4.41e-155 | COG4799@1\|root,COG4799@2\|Bacteria,1NRN3@1224\|Proteobacteria,1SA1V@1236\|Gammaproteobacteria,1X7S0@135614\|Xanthomonadales | 135614\|Xanthomonadales | I | Malonate decarboxylase | mdcE | 4.1.1.87 | ko:K13933 |
| accessory/8290/2/WP_107433696.1 | 522373.Smlt2725 | 1.98e-99 | COG0810@1\|root,COG0810@2\|Bacteria,1PKC7@1224\|Proteobacteria,1SFT0@1236\|Gammaproteobacteria,1X7D0@135614\|Xanthomonadales | 135614\|Xanthomonadales | M | TonB C terminal | - | - | - |
| accessory/8304/2/WP_132810695.1 | 522373.Smlt3799 | 1.34e-152 | COG0546@1\|root,COG0546@2\|Bacteria,1REXF@1224\|Proteobacteria,1S5AR@1236\|Gammaproteobacteria,1X67T@135614\|Xanthomonadales | 135614\|Xanthomonadales | S | hydrolase | idgB | 3.1.3.18 | ko:K01091 |
| accessory/8366/2/WP_049463140.1 | 391008.Smal_0990 | 1.72e-143 | COG3128@1\|root,COG3128@2\|Bacteria,1MUI7@1224\|Proteobacteria,1RQ0M@1236\|Gammaproteobacteria,1X383@135614\|Xanthomonadales | 135614\|Xanthomonadales | S | PkhD-type hydroxylase | - | - | ko:K07336 |
| accessory/8371/2/WP_132811132.1 | 522373.Smlt2379 | 4.54e-146 | COG0702@1\|root,COG0702@2\|Bacteria,1QR3E@1224\|Proteobacteria,1S37H@1236\|Gammaproteobacteria,1X7W7@135614\|Xanthomonadales | 135614\|Xanthomonadales | GM | epimerase | - | - | - |
| accessory/8492/2/WP_132810958.1 | 522373.Smlt4414 | 4.8e-149 | COG0346@1\|root,COG0346@2\|Bacteria,1PCRM@1224\|Proteobacteria,1SXWG@1236\|Gammaproteobacteria,1X6WW@135614\|Xanthomonadales | 135614\|Xanthomonadales | E | lactoylglutathione lyase activity | - | - | - |
| accessory/8551/2/WP_107433133.1 | 522373.Smlt3208 | 1.84e-155 | COG0625@1\|root,COG0625@2\|Bacteria,1REDI@1224\|Proteobacteria,1S13N@1236\|Gammaproteobacteria,1X4XE@135614\|Xanthomonadales | 135614\|Xanthomonadales | O | Belongs to the GST superfamily | gst5 | 2.5.1.18 | ko:K00799 |
| accessory/8749/2/WP_049398466.1 | 391008.Smal_2336 | 1.39e-150 | COG1335@1\|root,COG1335@2\|Bacteria,1MU5N@1224\|Proteobacteria,1RPGX@1236\|Gammaproteobacteria,1X30M@135614\|Xanthomonadales | 135614\|Xanthomonadales | Q | Hydrolase | - | - | - |
| accessory/8782/2/WP_005414428.1 | 522373.Smlt3946 | 5.22e-145 | COG2197@1\|root,COG2197@2\|Bacteria,1P4TD@1224\|Proteobacteria,1RYY9@1236\|Gammaproteobacteria,1X48T@135614\|Xanthomonadales | 135614\|Xanthomonadales | KT | LuxR family transcriptional regulator | - | - | - |
| accessory/8947/2/WP_110712507.1 | 522373.Smlt2721 | 1.57e-149 | COG1335@1\|root,COG1335@2\|Bacteria,1MU5N@1224\|Proteobacteria,1RPGX@1236\|Gammaproteobacteria,1X30M@135614\|Xanthomonadales | 135614\|Xanthomonadales | Q | Hydrolase | - | - | - |
| accessory/8999/2/WP_132810872.1 | 522373.Smlt4224 | 1.38e-138 | COG2197@1\|root,COG2197@2\|Bacteria,1RDKA@1224\|Proteobacteria,1S5A1@1236\|Gammaproteobacteria,1X4SE@135614\|Xanthomonadales | 135614\|Xanthomonadales | K | LuxR family transcriptional regulator | - | - | - |
| accessory/9034/2/WP_032130138.1 | 391008.Smal_0155 | 1.06e-142 | COG2197@1\|root,COG2197@2\|Bacteria,1MW84@1224\|Proteobacteria,1SPA2@1236\|Gammaproteobacteria,1XC3T@135614\|Xanthomonadales | 135614\|Xanthomonadales | K | luxR family | - | - | ko:K07687 |
| accessory/9207/2/WP_132810852.1 | 522373.Smlt4195 | 7.85e-151 | COG0625@1\|root,COG0625@2\|Bacteria,1RA4M@1224\|Proteobacteria,1S2K3@1236\|Gammaproteobacteria,1X9KM@135614\|Xanthomonadales | 135614\|Xanthomonadales | O | glutathione s-transferase | - | 2.5.1.18 | ko:K00799 |
| accessory/9216/2/WP_180844488.1 | 522373.Smlt1592 | 2.51e-126 | 2BZPC@1\|root,32RYC@2\|Bacteria,1N36B@1224\|Proteobacteria,1SVIW@1236\|Gammaproteobacteria,1X74Q@135614\|Xanthomonadales | 135614\|Xanthomonadales | - | - | - | - | - |
| accessory/9275/2/WP_132811080.1 | 522373.Smlt4679 | 3.45e-138 | COG0625@1\|root,COG0625@2\|Bacteria,1RA4M@1224\|Proteobacteria,1RYD2@1236\|Gammaproteobacteria,1XA6T@135614\|Xanthomonadales | 135614\|Xanthomonadales | O | glutathione s-transferase | gst8 | 2.5.1.18 | ko:K00799 |
| accessory/9387/2/WP_132809397.1 | 522373.Smlt1085 | 9.06e-129 | COG1028@1\|root,COG1028@2\|Bacteria,1RA3U@1224\|Proteobacteria,1RREY@1236\|Gammaproteobacteria,1X6EF@135614\|Xanthomonadales | 135614\|Xanthomonadales | IQ | Enoyl-(Acyl carrier protein) reductase | - | - | - |
| accessory/9772/2/WP_132810000.1 | 522373.Smlt2143 | 3.56e-117 | COG2201@1\|root,COG2201@2\|Bacteria,1RCWE@1224\|Proteobacteria,1S6II@1236\|Gammaproteobacteria,1X6PE@135614\|Xanthomonadales | 135614\|Xanthomonadales | NT | Chemotaxis response regulator containing a CheY-like receiver domain and a methylesterase domain | - | 3.1.1.61,3.5.1.44 | ko:K03412 |
| accessory/9796/2/WP_132809498.1 | 391008.Smal_1043 | 1.11e-125 | 2CDAN@1\|root,3134Z@2\|Bacteria,1RHTY@1224\|Proteobacteria,1S6GS@1236\|Gammaproteobacteria,1X7JC@135614\|Xanthomonadales | 135614\|Xanthomonadales | S | Gluconate 2-dehydrogenase subunit 3 | - | - | - |
| accessory/10489/2/WP_049399209.1 | 391008.Smal_3496 | 6.21e-122 | COG0346@1\|root,COG0346@2\|Bacteria,1RCYX@1224\|Proteobacteria,1RP3M@1236\|Gammaproteobacteria,1X69M@135614\|Xanthomonadales | 135614\|Xanthomonadales | E | Lactoylglutathione lyase | - | 4.4.1.5 | ko:K01759 |
| accessory/11407/2/WP_049399316.1 | 522373.Smlt2381 | 3.32e-91 | COG0745@1\|root,COG0745@2\|Bacteria,1RD6H@1224\|Proteobacteria,1S623@1236\|Gammaproteobacteria,1X7MU@135614\|Xanthomonadales | 135614\|Xanthomonadales | T | cheY-homologous receiver domain | - | - | - |
| accessory/12096/2/WP_049422820.1 | 522373.Smlt2920 | 1.43e-100 | COG0346@1\|root,COG0346@2\|Bacteria,1RGDW@1224\|Proteobacteria,1S8TY@1236\|Gammaproteobacteria | 1236\|Gammaproteobacteria | E | lactoylglutathione lyase activity | - | - | - |
| accessory/12292/2/WP_107431600.1 | 391008.Smal_3309 | 2.9e-85 | COG0848@1\|root,COG0848@2\|Bacteria,1RDJZ@1224\|Proteobacteria,1S3TA@1236\|Gammaproteobacteria,1X7PZ@135614\|Xanthomonadales | 135614\|Xanthomonadales | U | Biopolymer transport protein ExbD/TolR | - | - | ko:K03559 |
| accessory/12682/2/WP_019337750.1 | 522373.Smlt2711 | 1.47e-86 | COG0784@1\|root,COG0784@2\|Bacteria,1N70P@1224\|Proteobacteria,1SZ7B@1236\|Gammaproteobacteria,1XCTV@135614\|Xanthomonadales | 135614\|Xanthomonadales | T | cheY-homologous receiver domain | - | - | - |
| accessory/12968/2/WP_071229462.1 | 522373.Smlt1834 | 1.2e-83 | COG0346@1\|root,COG0346@2\|Bacteria,1N3K0@1224\|Proteobacteria,1T6TU@1236\|Gammaproteobacteria,1X7G9@135614\|Xanthomonadales | 135614\|Xanthomonadales | E | lactoylglutathione lyase activity | - | - | - |
| accessory/13693/2/WP_049423870.1 | 391008.Smal_2791 | 1.7e-81 | COG3832@1\|root,COG3832@2\|Bacteria,1N064@1224\|Proteobacteria,1SFE1@1236\|Gammaproteobacteria | 1236\|Gammaproteobacteria | J | glyoxalase III activity | - | - | - |
| accessory/13696/2/WP_132810831.1 | 522373.Smlt4143 | 1.02e-73 | COG0346@1\|root,COG0346@2\|Bacteria,1RH4X@1224\|Proteobacteria,1SAVS@1236\|Gammaproteobacteria,1XAWA@135614\|Xanthomonadales | 135614\|Xanthomonadales | E | Glyoxalase-like domain | - | - | - |
| accessory/15922/2/WP_006452007.1 | 391008.Smal_2413 | 7.54e-40 | COG1983@1\|root,COG1983@2\|Bacteria,1NAUA@1224\|Proteobacteria,1SD7E@1236\|Gammaproteobacteria,1X85M@135614\|Xanthomonadales | 135614\|Xanthomonadales | KT | Stress-responsive transcriptional regulator | - | - | - |

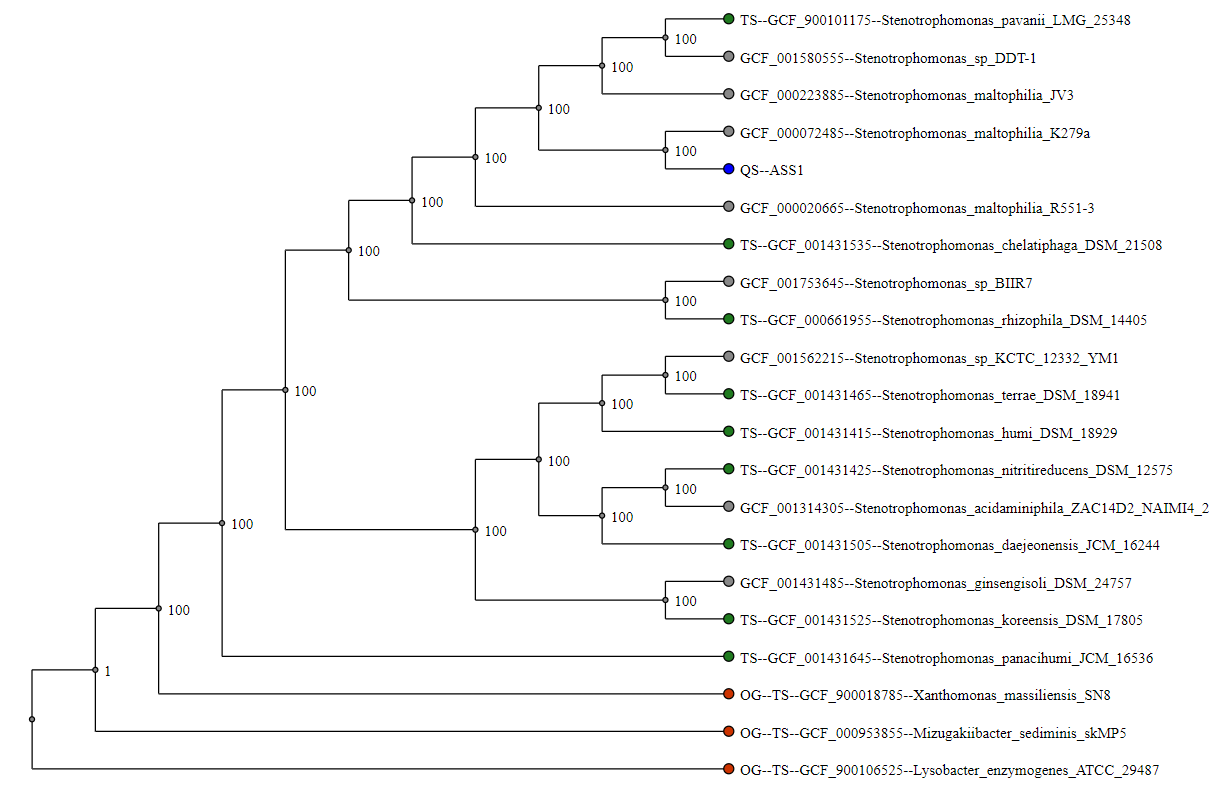

FigureS1: Concatenated Tree based on 89 MLST sequences from the genus Stenotrophomonas

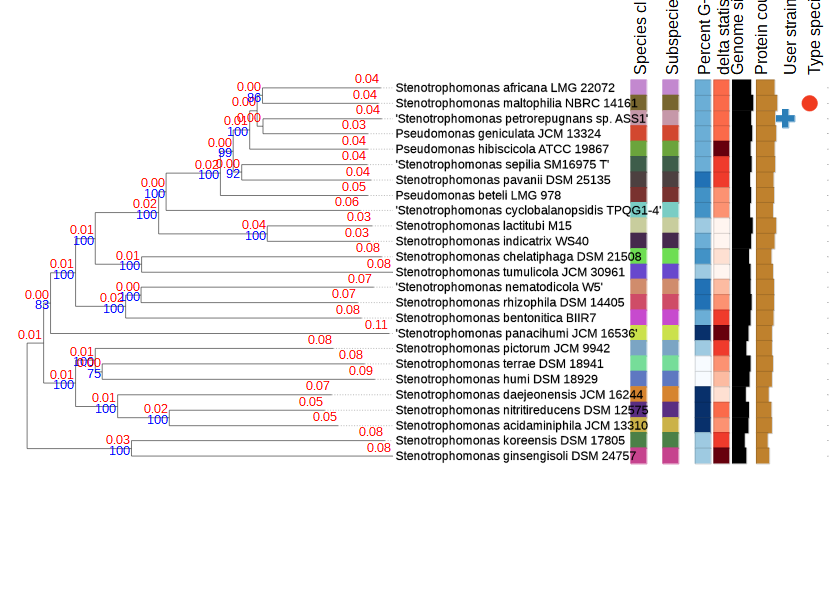

Figure S2: Phylogenetic tree was recovered from the TYGS database for the identification of typed genome. The tree was drawn based on the concatenated pan-genome of all typed Stenotrophomonas species and ASS1

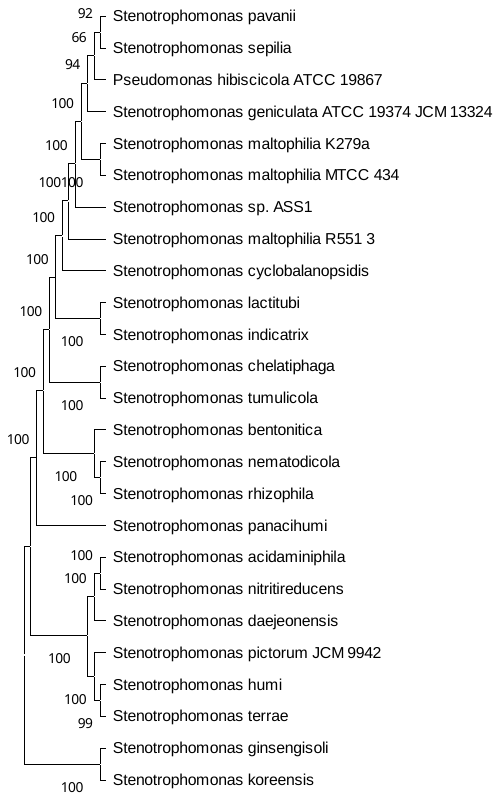

Figure S3: Phylogenetic tree drawn based on the complete genomes of the representative typed species and ***Stenotrophomonas sp* ASS1**

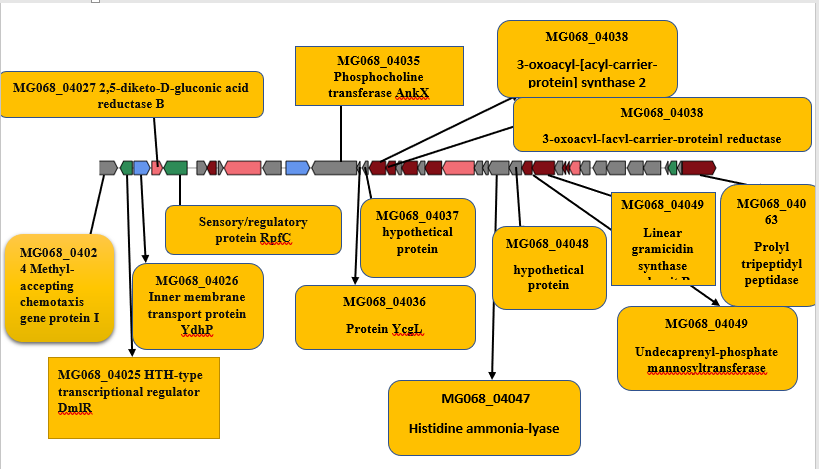

a. Arylpolyene cluster in ASS1

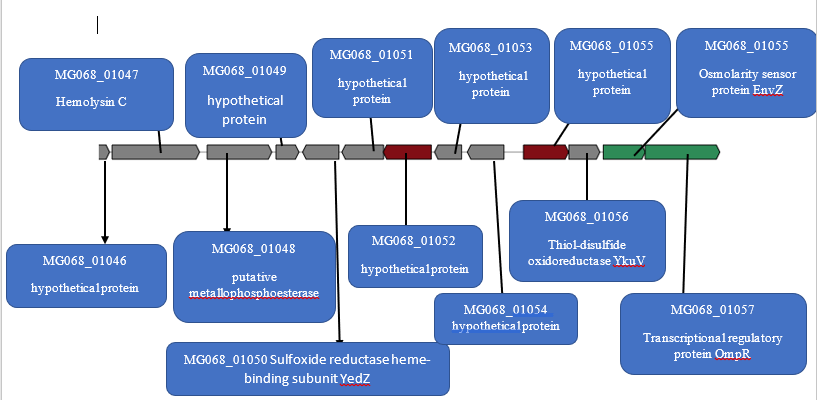

b. Bacteriocin cluster1 in ASS1

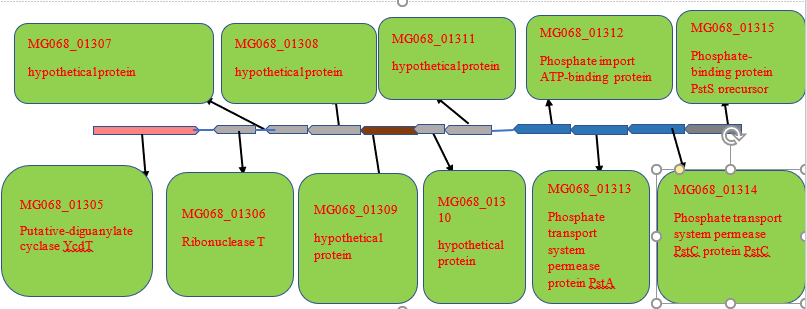

c. Bacteriocin Cluster II

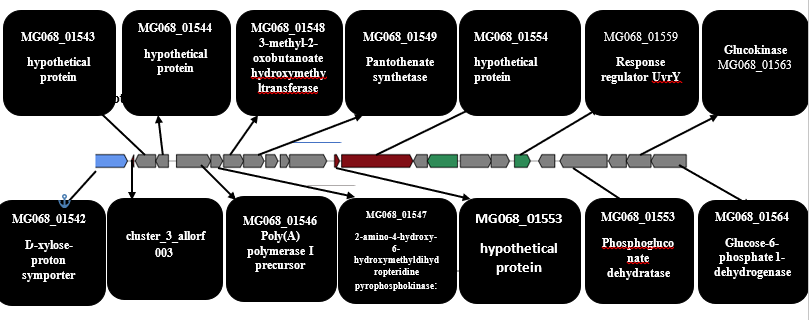

d. **Lantipeptide Cluster**

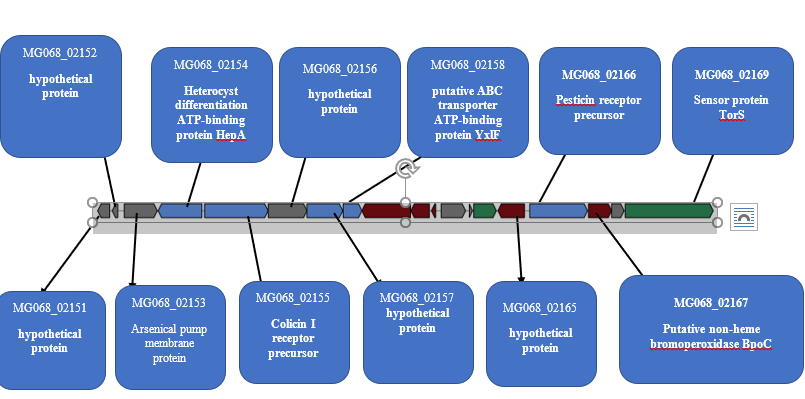

**Lassopeptide cluster**

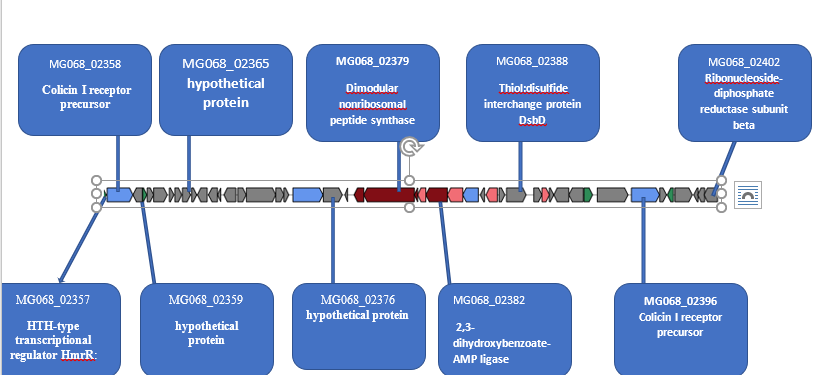

**Nrps Cluster associated with Hydrocarbon degradation**

**Figure S4**a-e Identified biosynthetic cluster in ASS1

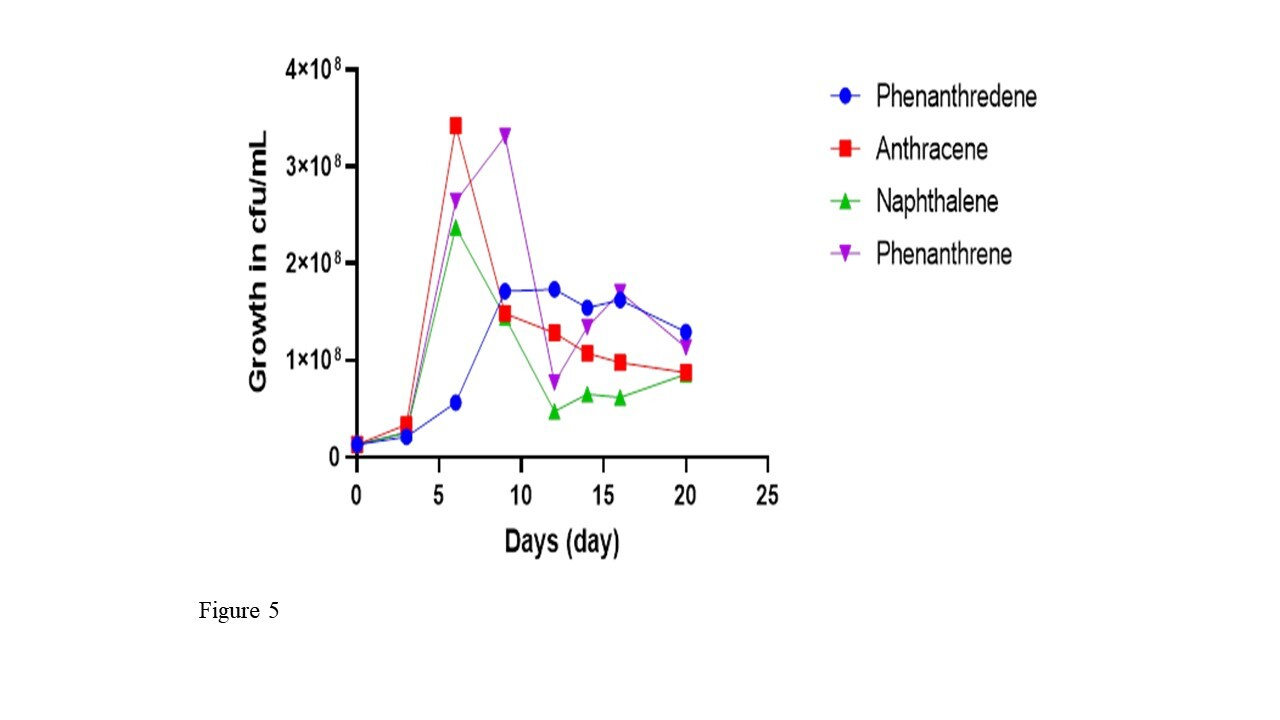

**Figure S5: ASS1 growth in different PAH as sole carbon source**

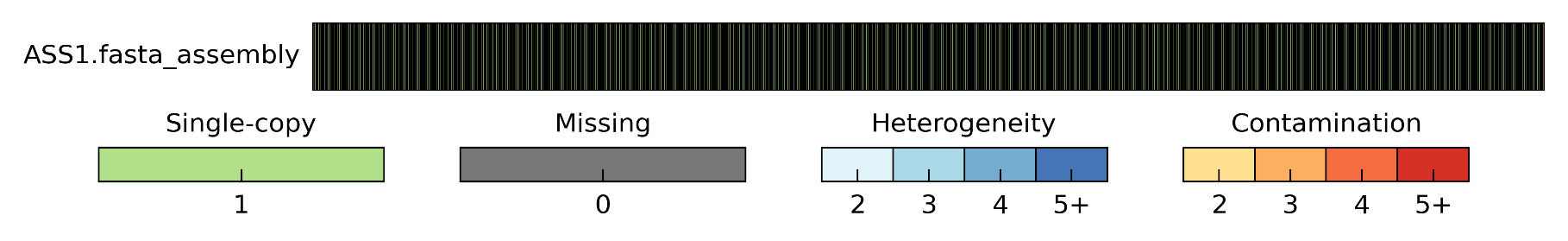

**Figure S6: Genome completeness file**
